# The finite state projection based Fisher information matrix approach to estimate information and optimize single-cell experiments

**DOI:** 10.1101/370205

**Authors:** Zachary Fox, Brian Munsky

## Abstract

Modern optical imaging experiments not only measure single-cell and single-molecule dynamics with high precision, but they can also perturb the cellular environment in myriad controlled and novel settings. Techniques, such as single-molecule fluorescence in-situ hybridization, microfluidics, and optogenetics, have opened the door to a large number of potential experiments, which begs the question of how best to choose the best possible experiment. The Fisher information matrix (FIM) estimates how well potential experiments will constrain model parameters and can be used to design optimal experiments. Here, we introduce the finite state projection (FSP) based FIM, which uses the formalism of the chemical master equation to derive and compute the FIM. The FSP-FIM makes no assumptions about the distribution shapes of single-cell data, and it does not require precise measurements of higher order moments of such distributions. We validate the FSP-FIM against well-known Fisher information results for the simple case of constitutive gene expression. We then use numerical simulations to demonstrate the use of the FSP-FIM to optimize the timing of single-cell experiments with more complex, non-Gaussian fluctuations. We validate optimal simulated experiments determined using the FSP-FIM with Monte-Carlo approaches and contrast these to experiment designs chosen by traditional analyses that assume Gaussian fluctuations or use the central limit theorem. By systematically designing experiments to use all of the measurable fluctuations, our method enables a key step to improve co-design of experiments and quantitative models.

**Author summary:** A main objective of quantitative modeling is to predict the behaviors of complex systems under varying conditions. In a biological context, stochastic fluctuations in expression levels among isogenic cell populations have required modeling efforts to incorporate and even rely upon stochasticity. At the same time, new experimental variables such as chemical induction and optogenetic control have created vast opportunities to probe and understand gene expression, even at single-molecule and single-cell precision. With many possible measurements or perturbations to choose from, researchers require sophisticated approaches to choose which experiment to perform next. In this work, we provide a new tool, the finite state projection based Fisher information matrix (FSP-FIM), which considers all cell-to-cell fluctuations measured in modern data sets, and can design optimal experiments under these conditions. Unlike previous approaches, the FSP-FIM does not make any assumptions about the shape of the distribution being measured. This new tool will allow experimentalists to optimally perturb systems to learn as much as possible about single-cell processes with a minimum of experimental cost or effort.

## Introduction

Recent labeling and imaging technologies have greatly increased capabilities to measure biological phenomena at the single-cell and single-molecule levels. When conducted under different conditions, single-cell experiments can probe processes for different spatial or temporal resolutions, for different population sizes, under different stimuli, at different times during a response, and for myriad other controllable or observable factors [1-7]. As these experiments have become more capable to precisely perturb or measure different biological species, they have also become more expensive, which imposes a limit on the number and type of experiments that can be conducted in any given study. Clearly, not all experiment designs provide the same information, and different experiments may be “optimal” to answer different questions about the system. However, the inherent diversity of modern experiments makes it difficult to intuit which experiments will be most informative and in which circumstances. Computational tools for model-driven experiment design could help to select more informative experiments, provided that existing tools can be adapted to overcome the unique challenges presented by single-cell data.

One model-driven approach to optimal experiment design is to use the *Fisher information matrix* (FIM), which describes the precision to which a model’s parameters can be estimated for any particular experiment [8-13]. To improve estimates of model parameters, the FIM can be used iteratively in a Bayesian framework by specifying maximally informative experimental conditions, collecting data under these conditions, using new data to constrain parameters, and using the newly constrained parameters to design the next round of experiments [9,12-15]. The formalism of the FIM for experiment design has been used to great effect in engineering disciplines, such as radar, astrophysics, and optics [16-18]. In principle, similar analyses could introduce a natural feedback in the co-design of single-cell experiments and discrete stochastic models, but for this to work, accurate analyses are needed to extract more meaning from the data and to provide better predictions about how biological systems will behave under new conditions.

Experimentally observed cell-to-cell variability has been well demonstrated to provide substantial quantitative insight to constrain and identify the mechanisms and parameters of gene regulation models [1-6,19-21]. Therefore, the FIM analysis for the optimal design of single-cell experiments should explicitly consider such single-cell variability. Standard FIM analyses assume continuous-valued observables with Gaussian-distributed *measurement* noise. However, in contrast to most classical engineering applications, the distributions of integer-valued RNA or protein levels across an isogenic cell population can be highly complex and subject to intrinsic and extrinsic variations, with nonlinear interactions that lead to multiple peaks and long tails [2,22-24]. Because the FIM is not computable for general discrete stochastic processes with non-Gaussian distributions, computational biologists have applied various approximations to estimate the FIM. A few recent biological studies use the Linear Noise Approximation [25] to treat single-cell distributions as Gaussian, which allows for the use of standard Fisher information analyses [8]. This approach, which we refer to as the LNA-FIM, should be valid for large numbers of molecules, but it is unlikely to be accurate for systems with high intrinsic noise corresponding to low gene, RNA, or protein counts. A different approach to estimate the FIM uses the central limit theorem (CLT) to approximate the sample mean and covariance to be jointly Gaussian and uses higher-order moments of the chemical master equation to estimate the likelihood of these moments [9]. This approach, which we refer to as the sample moments approach (SM-FIM), should be valid for large numbers of cells as can be collected in high-throughput experimental approaches, such as flow cytometry. However, when distributions have long asymmetric tails and sample sizes are limited, higher moments become very difficult to estimate and can lead to surprising model estimation errors [26]. Beyond these few Gaussian assumptions, there has been little work devoted to improve the design of time-varying single-cell experiments for systems with arbitrary probability distributions.

In this study, we introduce a formulation of the Fisher information for use with discrete stochastic models and data sets containing intrinsic variability that is measurable with single-biomolecule resolution. Our approach utilizes the finite state projection (FSP) approach [27] to solve the chemical master equation (CME) [25,28], and compute the likelihood of single-cell data given a discrete stochastic model [2,21,24]. The FSP solves for the probability distribution over discrete numbers of biomolecules to any arbitrary error tolerance. By utilizing the full probability distributions, as opposed to finite order or approximate moments of these distributions, our approach makes no assumptions and works well for distributions with multiple peaks or long tails.

In the next section, we introduce the FSP and derive the sensitivities of the FSP solution to small perturbations in parameters. Next, we derive the likelihood function and its local sensitivity for discrete stochastic models and discrete data. These allow us to formulate and compute the FSP-FIM. Next, we use a combination of analytical results and numerical simulations to verify the FSP-FIM for two common models of gene expression. Finally, we demonstrate how the FSP-FIM can be applied to design nontrivial experiments for a simulated system with nonlinear reaction rates.

### Chemical Master Equation and Finite State Projection

Stochastic gene expression can be modeled as a discrete state, continuous time Markov process, where different states 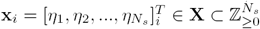 represent the *N*_*s*_ species of interest. In a biological context, the species *η* often correspond to gene configurations, RNA or protein abundances. Transitions to state **x**_*i*_ + *Ψ*_*v*_ from **x**_*i*_ occur with probabilities *w*_*v*_ (**x**_*i*_*, t*)*dt* in an infinitesimal time step of length *dt*, where *w*_*v*_ and *Ψ*_*v*_ are the propensity function and the stoichiometric vector corresponding to reaction *v* ∈ {1, 2*, …, N*_*r*_}. Using the propensity functions and stoichiometry vectors, one can describe the evolution of probability mass for each **x**_*i*_ using the chemical master equation (CME, [25,28]) given by:

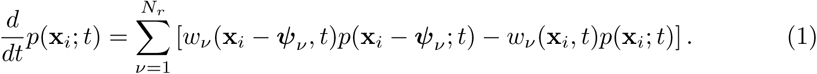

By enumerating all possible **x**_*i*_, one can define the probability mass vector as **p** = [*p*(**x**_1_; *t*)*, p*(**x**_2_; *t*)*, …*]^*T*^ and reformulate the CME in matrix form as 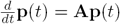.

Many systems described by the CME are not closed, i.e. the vector **p** has infinite dimension. In such cases, most states are extremely rare, and the sum of their corresponding probabilities is negligible. Thus, a natural approximation for the CME is to separate it into two exhaustive and disjoint sets, **X**_*J*_ and **X**_*J′*_, with **X**_*J*_ being a finite set and **X**_*J′*_ being a set of low probability states. Defining **p**_*J*_ (*t*) *≡ p*(**X**_*J*_; *t*), the CME can be reordered and written as:

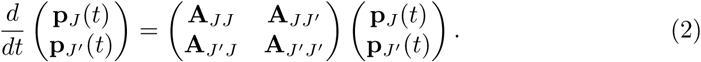

The finite state projection (FSP) approach [27], obtains an approximation of **p**_*J*_ (*t*) for finite times by replacing the set of states **X**_*J′*_ (*t*) with an absorbing sink state whose probability mass is *g*(*t*),

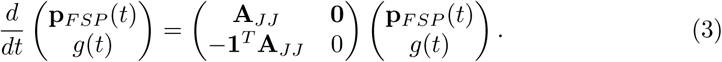

The FSP provides the exact total error of this approximation for all states in **X**_*J*_ and **X**_*J′*_ as:

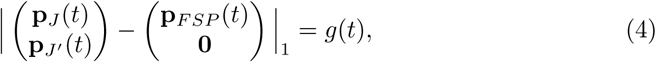

where the *|.|*_1_ denotes the absolute sum of the vector [24, 27]. The FSP solution is also guaranteed to be a lower bound on the true solution [24,27],

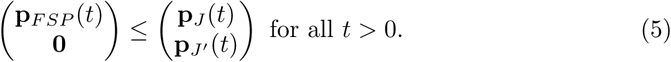

For simplicity, we will hereafter refer to the approximated states **p**_*F* *SP*_ (*t*) as **p**(*t*) and the corresponding matrix **A**_*JJ*_ as **A**. Next, we derive the likelihood function for FSP models and single-cell data.

### The FSP enables computation of the likelihood of single-cell data

A common task in single-cell analyses is to analyze snapshot measurements of independent cell populations, such as those collected using single-molecule fluorescent in-situ hybridization (smFISH) [22,23]. For such measurements, cells are fixed in the process of quantifying their RNA, and individual cells cannot be tracked over time. However, snapshots can be collected at different points in time to quantify a population’s response to changing conditions [2,29,30]. For such experiments, we assume that measurements at all time points {*t*_*k*_} are independent. The measured RNA counts for *N*_*s*_ different labeled species for each of *N*_*c*_ individual cells at time *t* can be collected into the data matrix 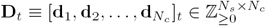. We define *L*(**D**; ***θ***) as the likelihood that all measured data 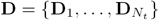 come from a model parameterized by ***θ*** = [*θ*_1_*, θ*_2_*, …*, *θ*_*k*_].

For FSP models, the likelihood and its logarithm for *N*_*c*_ measured cells can be written directly as:

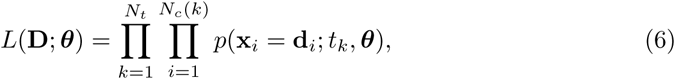

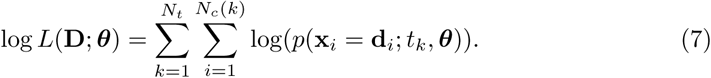

A common task in systems biology is to estimate parameters 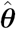 that maximize the likelihood that data could have come from a given model, and this form of the likelihood function has been used multiple times to estimate parameters from single-cell data [2, 6, 21, 24, 31, 32]. In addition to estimating parameters from data, the likelihood function can also be used to estimate the sensitivity of parameter estimates to sampling errors in the experimental measurements, which can in turn be used to design better experiments. In the following sections, we will use this fact to derive the FIM for FSP models.

### Derivation of the Fisher Information for FSP Models

The FIM, which describes the amount of information that can be expected by performing a particular experiment with *N*_*c*_ cells, is defined as

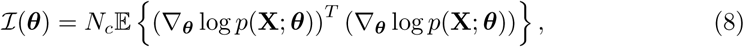

where the expectation is taken over *p*(**X**; ***θ***), corresponding to the density from which future (or hypothetical) data could be sampled. For FSP models, this density is the discrete distribution found by solving Eq. 3. Equation 8 is positive semi-definite and is additive for collections of independent observations [10]. The inverse of the FIM is known as the Cramèr-Rao bound (CRB), which provides a useful lower bound on the variance for any unbiased estimator of model parameters [11]. The notion of information stems from the fact that new experiments should increase the FIM, corresponding to additional knowledge about ***θ*** and a tighter CRB. More specifically, the well-known asymptotic normality of the maximum likelihood estimator (MLE) states that as the number of measurements *N*_*c*_ increases, the MLE estimates will converge in distribution to a multivariate normal probability density with a variance given by the CRB,

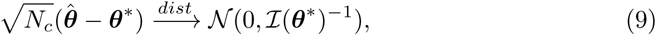

where 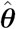 is the ***θ*** that maximizes Eq. 6 and ***θ***^*∗*^ are the “true” model parameters that produced the observed data [10,11]. Designing experiments to maximize a given metric of the FIM can be expected to provide a more accurate estimate of ***θ***, where different definitions of ‘accuracy’ (i.e., different vector norms for parameter errors) can be implemented through the choice of different FIM metrics.

To derive the FIM requires one must take the partial derivative of the log-likelihood (Eq. 7) with respect to the parameters ***θ***,

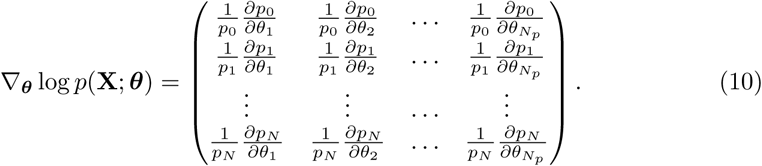

The expression *∇*_***θ***_*p*(**X**; ***θ***) is the *sensitivity matrix*, **S**, which has dimensions *N × N*_*θ*_, where *N* is the dimension of the CME or its FSP projection. As described in the Materials and Methods, we derive an equation similar to that presented in [33] to define the time evolution of the sensitivity for each state’s probability density, *p*(**x**_*l*_; ***θ***), to each parameter *θ*_*j*_. However, unlike previous analyses that rely on stochastic simulations and finite difference approaches, the FSP enables direct approximation of the sensitivities. Using the sensitivity matrix, the entries of the FIM can be computed as:

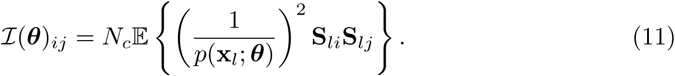

Taking the expectation over all *l* on (1*, N*) yields the elements of the FIM:

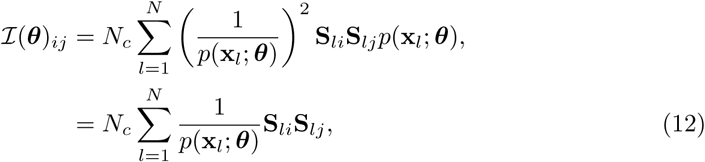

which quantifies Fisher information for the model evaluated at a single time point. For smFISH data, each time point is independent. If *N*_*c*_(*t*_*k*_) cells are measured at each *k*^th^ time point, the FIM is summed, and the total information is computed as:

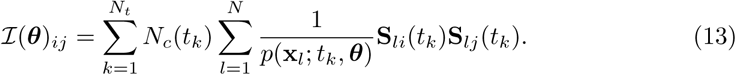

The Fisher information can be found using Eq. 13 for any model for which the FSP (Eq. 3) can be solved. This formulation explicitly quantifies how the number of cells and number of time points impact the information, and is easily extended to include other experiment design aspects such as the interval of successive measurements or changes in applied inputs, as we will demonstrate in the following sections. Because one is often interested in the relative sensitivity of parameters rather than the absolute sensitivity, a logarithmic parameterization of the FIM can easily be obtained from Eq. 13 by multiplying by the corresponding entries of ***θ*** (see supplemental information for full details),

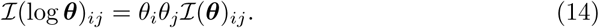

In the following sections, we will verify the FIM using several common models of gene expression, and demonstrate experiment designs using these approaches.

## Results

### The FSP-FIM captures the exact information for constitutive gene expression

To demonstrate and validate the FSP-FIM method, we begin with a simple birth and death model for constitutive gene expression as shown in Figure 1. This model, which has been fit to capture the variability for many housekeeping genes [1,20], consists of two reactions, corresponding to the constant transcription and first order decay of RNA,

**Fig 1.**
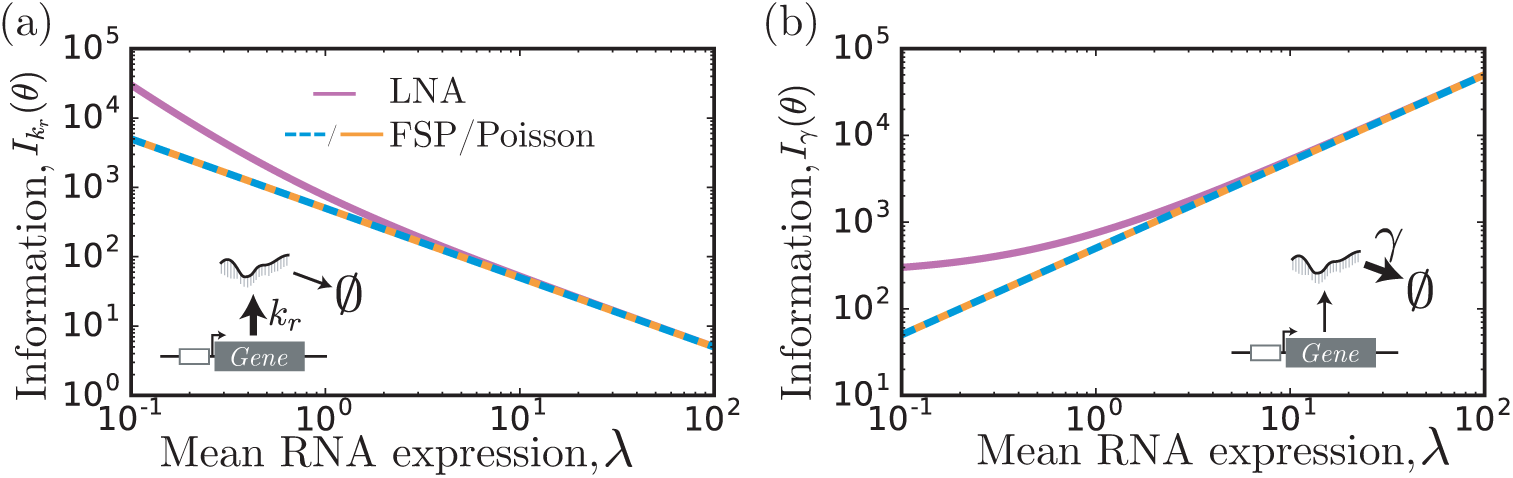
Fisher information for a model of birth and death. The Fisher information for the two model parameters *k*_*r*_ (a) and *γ* (b) for various values of the mean expression level, *λ*. The analytical form of the FIM for a Gaussian approximation and that computed using Eq. 37 (purple line) match to one another. The value computed using the FSP-FIM (blue) matches to the exact form of the analytical Poisson distribution (orange dashed). As *λ* becomes large, all four approaches are consistent.

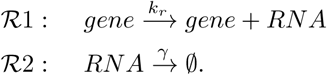

The production and degradation parameters are defined as ***θ*** = [*k*_*r*_*, γ*].

Given an initial condition of zero RNA for this process, the population of RNA at any later time is a random integer sampled from a Poisson distribution,

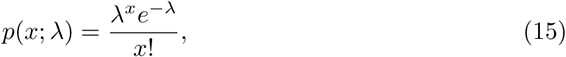

where *λ* is the time varying average population size,

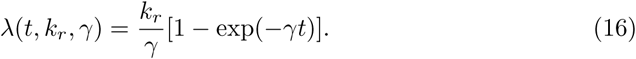

We have chosen the constitutive gene expression model to verify the FSP-FIM because the exact solution for the Fisher information for Poisson fluctuations can be derived in terms of *λ* as [10]:

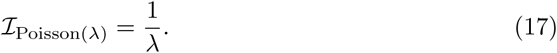

For convenience, the derivation of Eq. 17 is included in the supplementary text. Figure 1 shows the exact value of Fisher information (orange) versus the mean expression level for the two parameters *k*_*r*_ and *γ*. Figure 1 also shows that the FSP-FIM (blue) matches the exact solution for the information on both parameters at all expression levels, which verifies the FSP-FIM for this known analytical form.

Having demonstrated that the FSP-FIM matches to the exact solution, it is instructive to compare how well the previous LNA-FIM and SM-FIM estimates match to the exact FIM computation. For the Poisson distribution, the mean and variance are both equal to *λ*. Using this fact, the FIM can be approximated using the LNA-FIM for normal distributions (see Eq. 37 in the Materials and Methods). This expression, which is derived in the supplementary text, reduces to

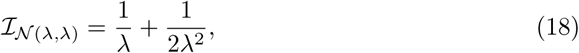

when both the mean and variance are *λ*. As *λ* becomes large, the Poisson distribution becomes well approximated by a normal distribution [11]. Equations 17-18 show that for this limit of large *λ*, the first term in Eq. 18 dominates, and ℐ_𝒩_ reduces to ℐ_Poisson_, yielding nearly equivalent values for the expected information. However at low mean expression *λ* ≤ 1, the strictly positive Poisson and the symmetric Gaussian distributions are less similar, and the Gaussian approximation predicts more information than is actually possible given the exact Poisson distribution. These trends are shown in Fig. 1, where the LNA-FIM approach only matches to the exact solution at high expression levels (compare orange and purple lines). For this example, the sample-moments based FIM (SM-FIM) is exact and matches to the analytical and FSP-FIM solutions at all expression levels [9].

### The FSP-FIM matches the simulated information for bursting gene expression

Next, we consider a slightly more complicated model of bursting gene expression, in which a single gene undergoes stochastic transitions between active and inactive states with rates *k*_on_ and *k*_off_. This switching model, which is depicted in Fig. 2(a), has been studied in detail [20,34-40], and it has been used to capture single-cell smFISH measurements in mammalian cells [30,37], yeast cells [2,36], and bacterial cells [29]. When active, the gene transcribes RNA with constant rate *k*_*r*_ and these RNA degrade in a first order reaction with rate *γ*. The four reactions of the system are:

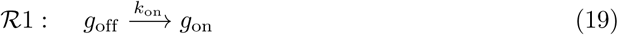

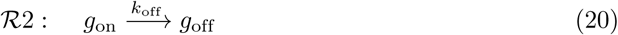

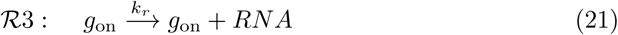

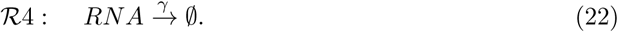

**Fig 2.**
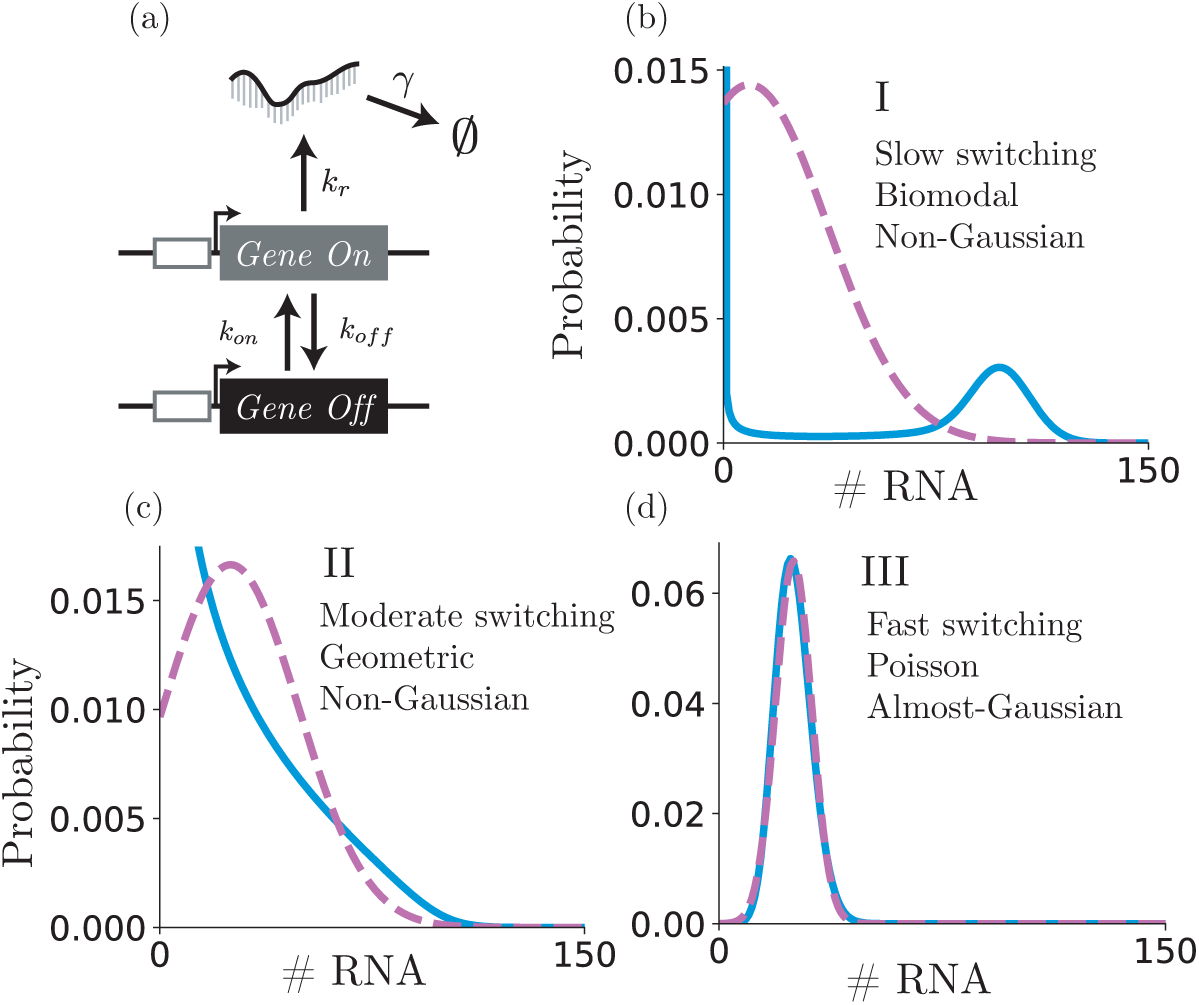
Bursting gene expression. (a) Schematic of the standard bursting gene expression model. Parameters are defined as given in the text to yield an “on” fraction of 0.25 and a mean expression of 25 mRNA per cell. (b) At slow switching rates, unique “on” and “off” modes are apparent, and distributions of molecule numbers are bimodal. (c) For intermediate switching rates, the distributions are geometric. (d) At high switching rates, the distributions are nearly Poisson (d). For each switch rate scale (labeled I, II, or III), the distribution of RNA computed with the FSP (blue) is compared to a Gaussian with the same mean and variance (purple).

For the examples below, we use the baseline parameters given by: *k*_on_ = 0.05*α* min^-1^, *k*_off_ = 0.15*α* min^-1^, *k*_*r*_ = 5.0 min^-1^, and *γ* = 0.05 min^-1^. In particular, the mRNA degradation rate, which sets the overall time-scale, was chosen to be representative of the average decay times (approximately 20 minutes) for mRNA in yeast [41].

For the bursting gene expression model, rescaling the transition rates *k*_on_ and *k*_off_ by a common factor does not affect the mean expression level, because the fraction of time spent in the active state remains unchanged. This fraction can be written

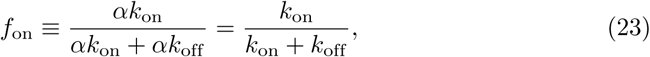

and is the same for any *α >* 0. For the parameters given above, the average expression at steady state is given by *k*_*r*_*f*_on_*/γ* = 25. However, rescaling the transition rates does change the shape of the distribution as shown in Fig. 2(b-d)[20]. When switching is slow, the gene stays in the “on” and “off” states long enough to observe individual high and low peaks corresponding to the “on” and “off” states, as in shown in Fig. 2(b). However, for intermediate switching rates, the gene does not spend enough time in the “off” state for bursts to decay or enough time in the “on” state for large populations to accumulate (see Fig. 2(c)). At fast switching rates the “on” and “off” states come to a fast quasi-equilibrium, and the time-averaged system approaches a Poisson process, where the effective production rate is *k*_*r*_*f*_on_. For the bursting gene expression model, all moments of the distributions can be computed exactly from Eq. 35 in the Materials and Methods section, even when the RNA distributions are highly non-Gaussian [42].

Since the previous example has already verified the accuracy of the FSP-FIM when the expression has a Poisson distribution, we now verify the FSP-FIM for the slow switching case in which the distribution is bimodal (*α* = 0.1). To our knowledge an exact FIM solution is not known for the bursting gene expression model, so we evaluate the different FIM approximations by finding the sampling distribution of the MLE, and we compare the covariance of this distribution to the inverse of the FIM [11]. To do this, we sample from *p*(**X**; *t*, ***θ***^*∗*^) under reference parameter set ***θ***^*∗*^ to generate 200 simulated data sets, each with independent RNA measurements for 1,000 cells. We then allow *k*_off_ and *k*_*r*_ to be free parameters, and we find 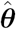 for each of the 200 data sets. Figure 3 compares the 95% confidence intervals found by taking the inverse of the FIM and through MLE estimation using simulated data for the FSP likelihood (Eq. 6) shown in Fig. 3(a), the LNA-based likelihood (Eq. 36 in the Methods section) shown in Fig. 3(b), and the SM-based likelihood (Eq. 36 in the Methods section, Supplementary Eq. 10) shown in Fig. 3(c). Figure 3(a) shows that the CRB predicted by the FSP-FIM matches almost perfectly to the confidence intervals determined by MLE analysis of independent data sets. Figure S3 (left column) shows that this estimate is consistently accurate over multiple different experiment designs. In contrast, the LNA-FIM dramatically overestimates the information and predicts confidence intervals that are much smaller than are actually possible (Figs. 3(b) and S3, center column). The SM-FIM does a better job than the LNA in that it matches the MLE analysis for some experimental conditions (Fig. 3(c)) but much less well for other conditions (Fig. S3, right column). We note that the three different FIM estimates yield different principle directions and different magnitudes for parameter uncertainty (Fig. 3(d)), but in all cases the FSP-MLE matches to the FSP-FIM and results in the tightest MLE estimation.

**Fig 3.**
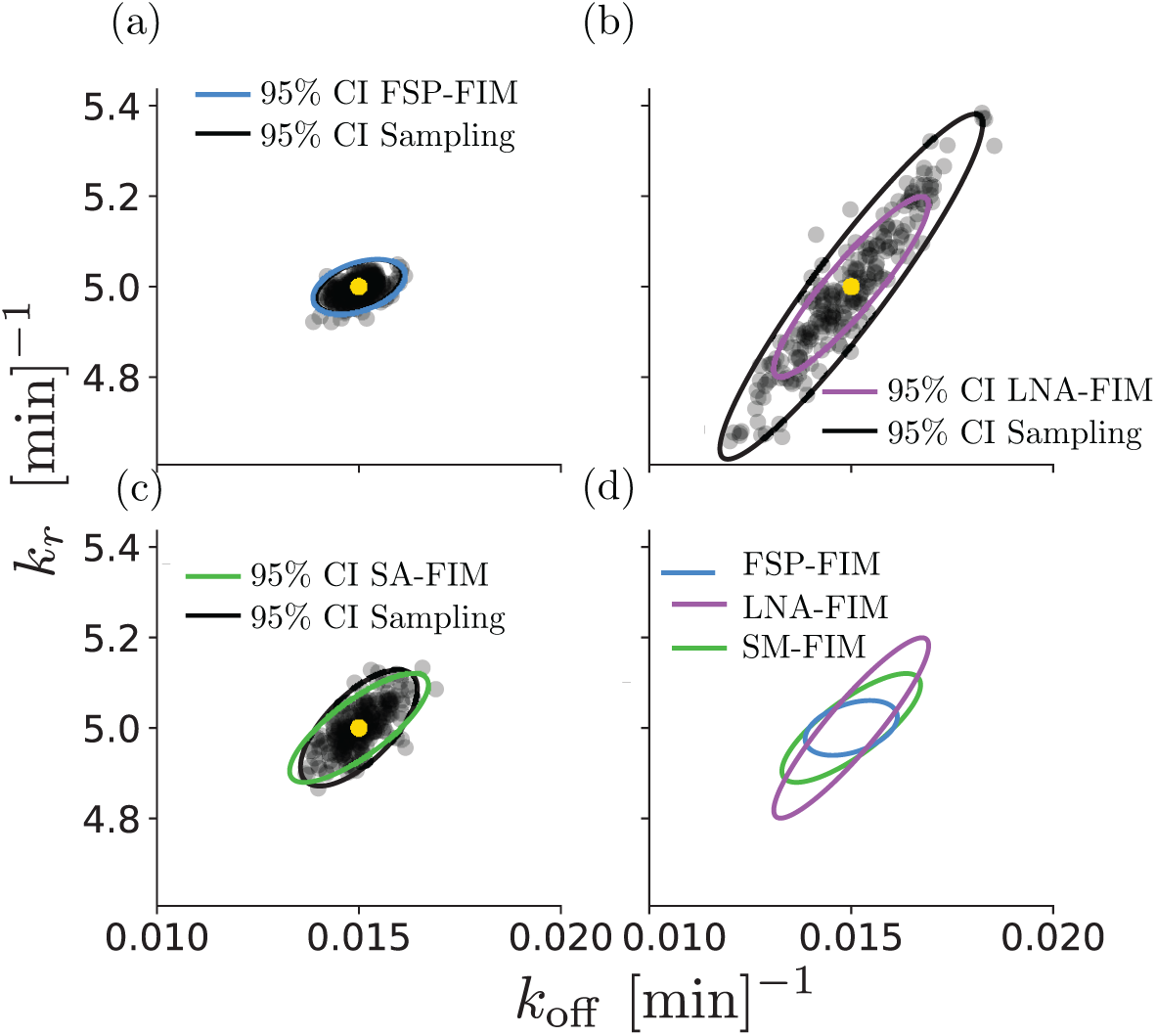
Verification of the FSP-FIM for models with non-Gaussian distributions. The inverse of the FIM is a lower bound on the variance of the MLE estimator. Here, we simulate 200 data sets with 1,000 cells in each data set. We then find the MLE 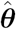 (scatter plots) for each, and compare the covariance of these samples to the inverse of the FIM for the (a) FSP-, (b) LNA-, and (c) SM-FIM approaches. Panel (d) shows the FIM matrices for all approximations on the same axes. Simulated data were generated using the parameters given in the main text and at 10 time points evenly distributed between 0 and 200 minutes.

Having verified the FSP-FIM for the bursting gene expression model with multiple parameter sets, we next explore how the information changes as a function of the system parameters. Figure 4 shows the determinant of the FIM (also known as the D-optimality or information density) for the bursting gene expression model as a function of the switch rate scaling factor, *α*, using the LNA-FIM (purple), SM-FIM (green) and FSP-FIM (blue) approximations. In the limit of fast switching (i.e. *α* → *∞*), the expected information converges to nearly the same value for all approaches, as expected for a Poisson distribution with high effective population size of *λ* = 25 RNA. However, in the non-Gaussian regimes with slow switch rates, the LNA-FIM over-estimates and SM-FIM under-estimates the information compared to the verified FSP-FIM approach. We note that these differences arise despite the fact that the bursting gene expression model has linear propensity functions, which allows for closed and exact computation of the statistical moments.

**Fig 4.**
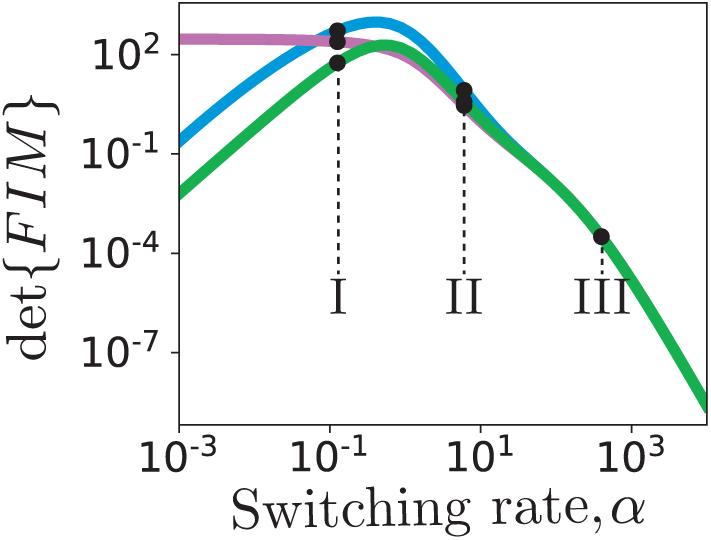
FIM analysis of the bursting gene model. The determinant FIM for the LNA-FIM (purple), FSP-FIM (blue), and SM-FIM (green) as a function of the gene switching rate scale, *α*. Labels I, II, III correspond to the switch rates for which distributions are plotted in Figs. 2(a-c). Parameters are given in the main text and data are assumed to be collected at 10 equally separated time points between 0 and 200 minutes.

### The FSP-FIM Can Design More Informative Single-Cell Experiments

Next, having verified the FSP-FIM for its ability to accurately estimate the FIM for different parameter sets, we explore the use of the FSP-FIM to design experiments that maximize information. Specifically, we will use classical FIM-based experiment design approaches to choose single-cell experiments first for the bursting gene expression model above, and then for a nonlinear toggle model for which moments can no longer be computed exactly. We consider two different metrics of the FIM, which are frequently used in model-driven experiment design [9,12]. The first of these is E-optimality presented in the main figures), which corresponds to the smallest eigenvalue of the FIM. By finding the experiment which maximizes this eigenvalue, the information is increased in the principle direction of parameter space in which the least information is known (i.e. the parameter uncertainty is highest). The second FIM criteria is D-optimality (presented in supplemental figures), which corresponds to the determinant of the FIM. By maximizing the determinant of the FIM over the experiment design space, one finds an experiment which minimizes the volume of the uncertainty in parameter space. We note that many other experimental design criteria are possible, and the choice of criteria depends on what one desires to learn about the system.

#### Optimizing the sampling rate for bursting gene expression

Our first demonstration of FSP-FIM based experiment design is to select the optimal single-cell sampling period with which to identify the parameters of the bursting gene expression model. For this, we have chosen to analyze E-optimality criteria, which seeks to maximize the smallest eigenvalue of the FIM. We consider a potential experiment design space consisting of 60 logarithmically distributed sampling periods Δ*t* from 2 *×* 10^-2^ minutes and 7 *×* 10^2^ minutes. For each sampling period, a total of five evenly spaced temporal measurements would be taken. Figure 5(a) compares the information expected versus the sampling period using the different FIM approximations: LNA-FIM (purple), SM-FIM (green) and FSP-FIM (blue). For each potential experiment, we then simulate 200 data sets for 1,000 cells each by sampling *p*(**X**; *t*, ***θ***^*∗*^), use Eq. 7 to find the MLE parameter estimate for each data set, and then compute the covariance matrix from the MLE parameter sets. This covariance matrix is inverted, and its minimum eigenvalues are depicted as orange triangles in Fig. 5(a). Figure 5(b) also shows a scatterplot to compare the relationship between the MLE-observed information and the predicted information for all FIM approaches. Once again, the FSP-FIM consistently matches the observed E-optimality at all experimental conditions. However, the LNA approach is much less consistent, sometimes over-estimating and sometimes under-estimating the real information for the different experimental conditions. The SM-FIM consistently underestimates the true information for this example, although it is not clear if this trend would hold for all sets of parameters and experimental conditions.

**Fig 5.**
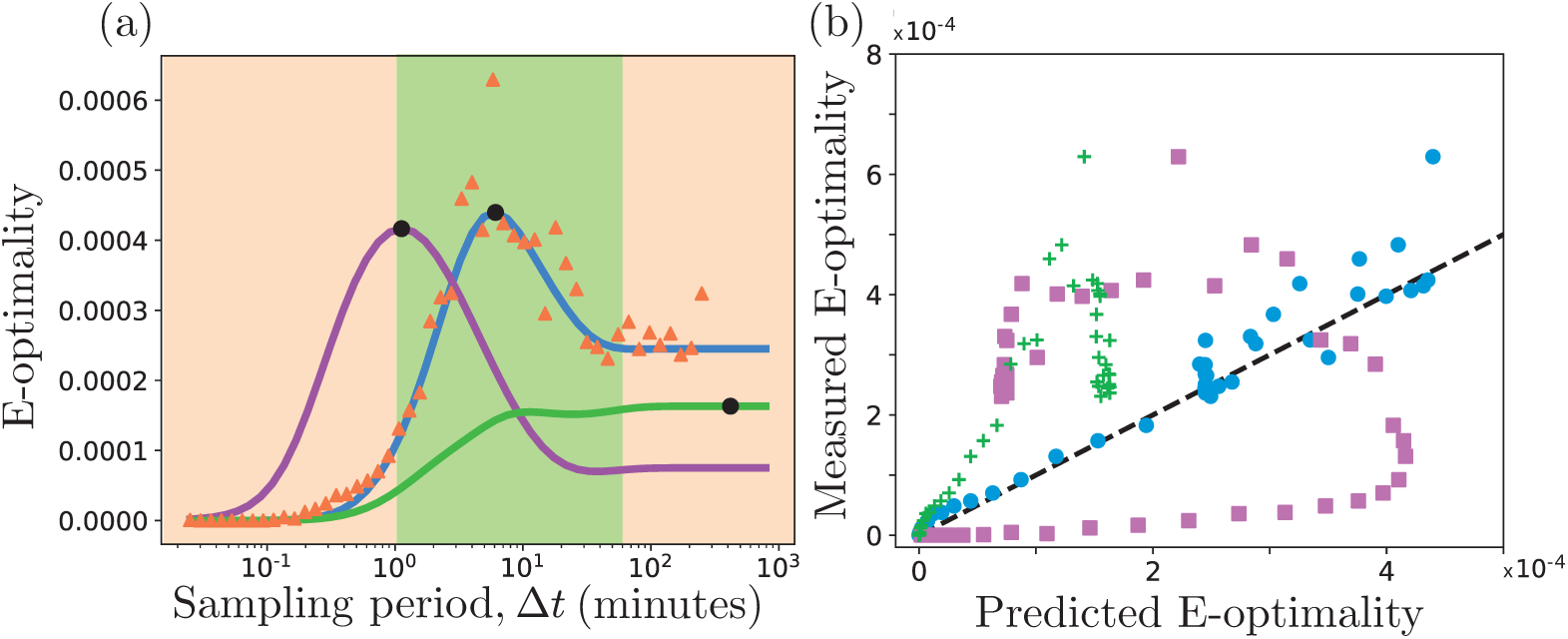
Designing experiments with the FSP-FIM. (a) E-optimality (i.e., smallest eigenvalue of the FIM) for the standard bursting gene expression model versus sampling period, Δ*t*, using FSP-FIM (blue), LNA-FIM (purple), and SM-FIM. Maximizing E-optimality corresponds to minimizing variance in the in the most variable direction of parameter space. The orange triangles show MLE-based confirmation of the E-optimality, using 200 simulated data sets for each sampling period. The green shaded region represents experiments that are feasible using smFISH, from minute resolution [2] to hour resolution [29] (b) Comparison of the FSP-FIM (x-axis) versus the observed information (y-axis) for various sampling periods using the FSP-FIM (blue circles), LNA-FIM (purple squares), and SM-FIM (green crosses). Kinetic parameters are given in the main text.

From Fig. 5(a), it is clear that the amount of expected information depends strongly on the sampling period. When the sampling period is much longer than the characteristic time to reach the steady state distribution (Δ*t* ≫1*/γ*), the information does not change because all snapshots are already close to steady state. When the sampling period is too short (Δ*t* ≪ 1*/γ*), there is insufficient time for the distributions to change and the information tends to zero. Despite conserving these trends, the three different FIM analyses result in substantially different optimal experiments for the E-optimality design criteria. Using the FSP-FIM, the optimal experiment is Δ*t* = 6.1 minutes, which we verified using the MLE sampling approach (compare orange triangles and blue line in Fig. 5(a)). This optimal design is well-aligned with smFISH experimental technique, which can capture cell populations with one minute resolution [2] to one hour resolution [29]. However, the LNA-FIM selects a much faster sampling period of Δ*t* = 1.1 minutes, and the SM-FIM selects a much slower sampling period of Δ*t* = 420 minutes. Thus, the FSP-FIM not only provides more information compared to moments-based approaches, but it also provides a better estimate of the expected information. In turn, these improved estimates can help to avoid potentially misleading experiments and select optimal designs.

#### The FSP-FIM accurately estimates information for systems with nonlinearities and bimodal responses

To demonstrate the utility of the FSP-FIM approach for models with nonlinear reaction propensities and multiple species, we turn to the toggle model first introduced by Gardner et al [43], with a stochastic formulation by Tian and Burrage [44]. Figure 6(a) shows a schematic of the toggle model, which consists of two mutually repressing proteins, *x* ≡ LacI and *y* ≡ *λ*cI, where the production of each species depends non-linearly on the concentration of its competitor. The reactions in the toggle model can be written

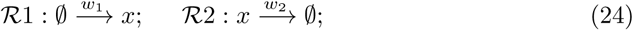

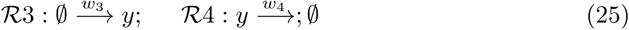

where

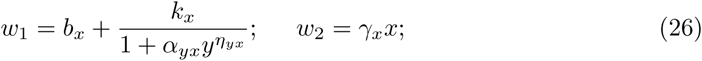

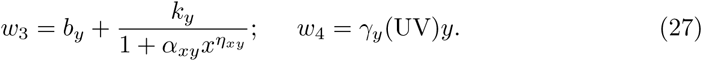

**Fig 6.**
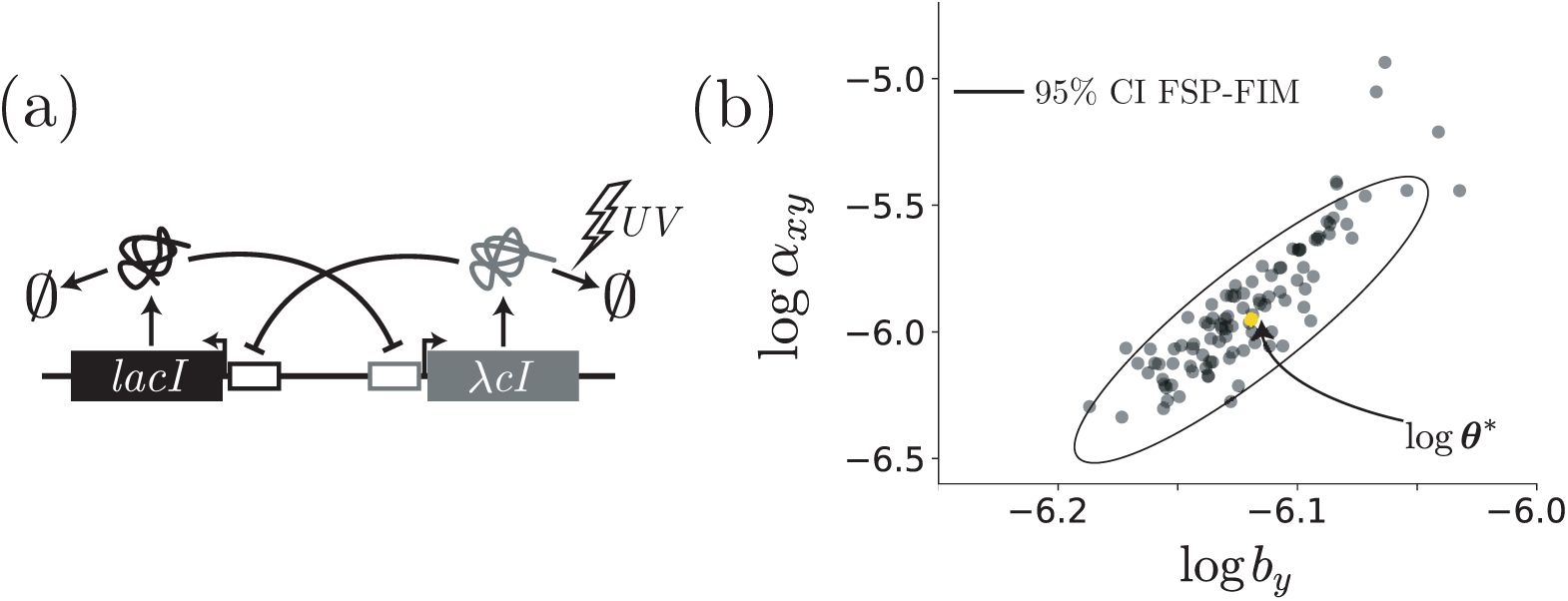
Validation of a toggle model. (a) Model schematic of the two genes, *lacI* and *λ*cI, which are mutually repressing [43]. Degradation of *λ*cI is controlled by UV radiation. (b) Verification of the FSP-FIM (black ellipse) for 200 MLE estimates of 1,000 cells each (black dots) for two free model parameters, *α*_*xy*_ and *b*_*y*_.

In this formulation, we have assumed that the degradation of *λcI* is controlled by an ultraviolet (UV) radiation through the light-induced circuit described by Kobayashi et al [45]. Similar to [46], we assume that the UV level affects the degradation of *λ*cI according to the function:

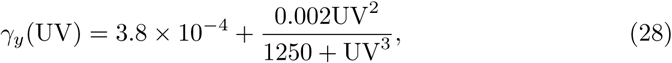

where the minimum degradation rate has been chosen to match dilution due to the *E. coli* half life of 30 min [46]. The remaining parameters used for this example are given by ***θ***^*∗*^ in Table 1. The system’s initial condition at *t* = 0 is assumed to be the equilibrium distribution when no UV is applied. For this biological system and these parameters, different levels of UV radiation will give rise to different dynamics. At low levels of radiation, switching to the high LacI state is rare, and the distribution tends to have a single peak. At intermediate levels of radiation, switching between low and high levels of LacI expression is possible, and LacI distributions may be bimodal. Finally, at high levels of radiation, the system very quickly switches into the high LacI state.

**Table 1.** Parameters for the toggle model. ***θ***^*∗*^ is the “true” parameter set from which data is generated, and 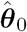 is the MLE parameter set fit to a baseline data set generated assuming 0 UV (see Fig. S5 for further discussion). Here, *N* is used to denote the units of single-molecules.

| | $\theta^*$ | $\hat{\theta}_0$ | units |
| --- | --- | --- | --- |
| $b_y$ | $6.80 \times 10^{-5}$ | $9.86 \times 10^{-4}$ | $s^{-1}$ |
| $b_x$ | $2.20 \times 10^{-3}$ | $3.19 \times 10^{-3}$ | $s^{-1}$ |
| $k_y$ | $1.60 \times 10^{-2}$ | $1.60 \times 10^{-2}$ | $s^{-1}$ |
| $k_x$ | $1.70 \times 10^{-2}$ | $2.50 \times 10^{-2}$ | $s^{-1}$ |
| $\alpha_{xy}$ | $6.10 \times 10^{-3}$ | $8.28 \times 10^{-3}$ | $N^{-\eta_{xy}}$ |
| $\alpha_{yx}$ | $2.60 \times 10^{-3}$ | $2.46 \times 10^{-3}$ | $N^{-\eta_{xy}}$ |
| $\eta_{xy}$ | 2.10 | 2.10 | - |
| $\eta_{yx}$ | 3.00 | 3.00 | - |
| $\gamma_x$ | $3.80 \times 10^{-4}$ | $5.57 \times 10^{-4}$ | $N^{-1}s^{-1}$ |

Because this model has complex nonlinear propensity functions, the statistical moments cannot be calculated in closed form, and the LNA-FIM and SM-FIM estimates are no longer expected to provide accurate estimates for information or optimal experiment designs. In contrast, the FSP analysis remains unchanged, and the FSP-FIM can be computed exactly as above. As before, we verify the FSP-FIM for this nonlinear case using a set of 200 simulated data sets measured at 1 hr, 4 hr, and 8 hr, each with 1,000 cells, and we found MLE parameter estimates 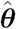 for each simulated data set. Figure 7(a) shows this verification in a simple case with two free parameters, *b*_*y*_ and *α*_*xy*_, and Fig. S4 shows the verification where all parameters free except for Hill coefficients *η*_*xy*_ and *η*_*yx*_. In this and all subsequent analysis of the toggle model, we have used the logarithmic parameterization of the FIM (Eq. 14).

**Fig 7.**
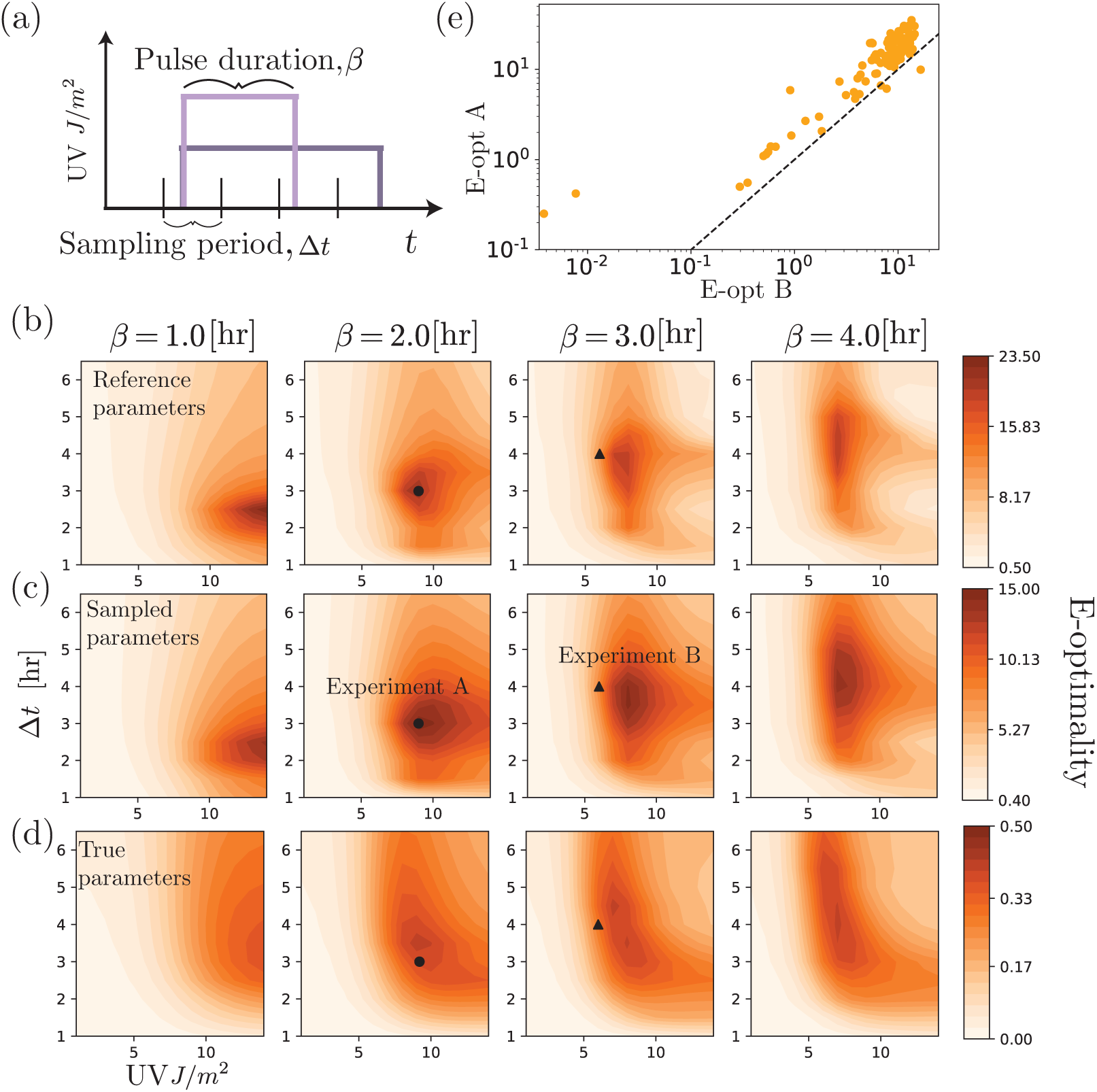
Experiment design for the nonlinear genetic toggle model. (a) Degradation rate of *λ*cI is controlled by UV as shown in Fig. 6(a). The magnitude and duration (*β*) of UV exposure are free experiment design parameters, along with the time between measurements Δ*t*. (b) E-optimality (the smallest eigenvalue of the FIM) versus the 3-dimensional experiment design space, where the FIM is computed using (b) the reference parameter set, (c) by averaging the E-optimality over 100 unique parameter sets and (d) using the “true” parameter values. The black circle is the optimal design chosen according to (c). The black triangle denotes a nearby, but less informative, experiment. (e) For the experiments corresponding to the black circle and triangle in (b-d), E-optimality values are shown for each sampled parameter set.

Next, we aim to design more complex experiments for the toggle model described above. We consider an experiment design space where the measurement sampling period (Δ*t*), pulse duration (*β*), and pulse magnitude (UV) can all be changed, as illustrated in Fig. 7(a). Each pulse of UV starts at *t* = 1 hr. We then compute the FSP-FIM for each experiment {UV*, β*, Δ*t*}.

To capture the more realistic situation where parameters are unknown prior to experimentation, we next explore how parameter uncertainty affects the estimation of the FIM and the design of optimal experiments. To begin, we assume that parameters have been partially estimated from a simple initial experiment corresponding to measurements of the unperturbed steady state at zero UV input to the system. In practice, similar preliminary parameter estimates could be acquired from literature, from previous less-optimized experiments, or by comparison to related pathways or organisms. For our analysis, the prior estimate for parameters is described by a multivariate lognormal distribution with a geometric mean of 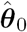 given in Table 1 and covariances given in Table S1. Parameters sampled from this distribution are substantially different from the “true” parameter, *θ*^*∗*^, which is also shown in Table 1. Figure 7(b) shows the E-optimality criteria for parameter set 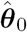 as a function of the experiment design parameters {UV*, β*, Δ*t*}. Next, we sampled 100 random sets of parameters from the prior distribution (Fig. S5), and we computed the E-optimality for each set. Figure 7(c) presents expected information versus experiment design averaged over these 100 parameter sets. For comparison, Fig. 7(d) shows the information versus experiment designs if one had exact knowledge of the true parameters.

From Figs. 7(b-d), we observe that relative estimates of the FIM remain consistent despite substantial changes to the parameters from which the FIM is computed. To explore this observation more closely, we selected the experiment that maximizes the averaged E-optimality in Fig. 7(c). This experiment is denoted by a black circle in Figs. 7(b-d), and we compare it to another similar experiment design, shown by the black triangle in Fig. 7(b-d). Figure S6 shows the expected parameter uncertainty for these two designs and shows that the optimal experiment reduces variance in some parameter directions by more than an order of magnitude compared to the sub-optimal experiment. To explore how different parameters change the ranking of these two experiments, we analyze the ranking of Experiment A and Experiment B not only based on their average E-optimality value as in Fig. 7(c), but at each of 100 random parameter combinations. Figure 7(e) shows that for 97 of the 100 parameter samples, the relative ranking of the experiments is consistent, even though the absolute value of the E-optimality criteria varies over several orders of magnitude.

We next seek to understand how optimal experiments depend on one’s plans to perform multiple experiments. The “single experiment” in Table 2 refers to designing a single experiment, *ε*_1_, to maximize the expected FIM design criteria, such as finding the maximal combination in Fig. 7(c). The “dual greedy” approach also chooses the same *ε*1 and then seeks to find the most complementary additional experiment, *ε*_2_, to maximize the overall FIM design criteria. Finally, the “dual simultaneous” search finds the optimal combination of any two possible experiments, 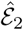 and 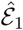 to maximize the design criteria. Since the optimal choice for 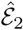 and 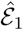 can admit the other choices, it must yield at least as high a design criteria as *ε*_1_ and *ε*_2_. By comparing the three design strategies for the current toggle model, we find indeed that the simultaneous approach discovers a substantially more informative experiment than does the greedy approach. In other words, the overall expected value of an experiment, can depend not only on the current parameter values, but also upon which other experiments one intends to conduct. If one has plans to do multiple experiments, it may be better to consider the potential information from all experiments as a whole rather than to design each experiment one at a time.

**Table 2.** Comparing sequential experiment design approaches.

|  | Single experiment | Dual greedy | Dual simultaneous |
| --- | --- | --- | --- |
| $\begin{Bmatrix} \beta \\ \Delta \\ \text{UV} \end{Bmatrix}$ | $\begin{Bmatrix} 2 \text{ hr} \\ 3 \text{ hr} \\ 9 \text{ J/m}^2 \end{Bmatrix}$ | $\begin{Bmatrix} 4 \text{ hr} \\ 5.5 \text{ hr} \\ 14 \text{ J/m}^2 \end{Bmatrix}$ | $\begin{Bmatrix} 1 \text{ hr} \\ 2.5 \text{ hr} \\ 9 \text{ J/m}^2 \end{Bmatrix}, \begin{Bmatrix} 4 \text{ hr} \\ 2.5 \text{ hr} \\ 13 \text{ J/m}^2 \end{Bmatrix}$ |
| E-opt | 14.9 | 32.0 | 36.8 |

## Discussion

Fluctuations in biological systems complicate modeling by introducing substantial variability in gene expression among individual cells within a homogeneous population. This variability contains valuable and quantifiable insights [20], but data with multiple peaks and long tails, such as those collected using smFISH, are often poorly described by modeling approaches that only make use of low-order moments of such distributions [26]. The FSP approach [27] has previously been used to identify and predict gene expression dynamics for complex and rich single-molecule, single-cell data [2,29,30]. In this work, we have developed the FSP-based Fisher information matrix, which extends the FSP analysis to allow rigorous design of experiments that are optimally informative about the model’s parameters.

The FSP-FIM uses a novel sensitivity analysis, which requires solving a system of ODEs that is twice the size of the FSP dimension for each parameter, and therefore should be computationally tractable for any problem to which the FSP can be applied. The local sensitivity of each parameter is independent of the other parameters, so the computation is easily parallelized among multiple processors. We verified that the FSP-FIM approach matches the information for the constitutive gene expression model, whose response follows a Poisson distribution (Fig. 1), and for which the FIM can be computed exactly. The FSP-FIM also matches to classical FIM approaches that assume normally distributed data (LNA-FIM) or very large data sets (SM-FIM) in the limiting case when the data distributions are close to being Gaussian (Figs. 1-4). For systems where data is not Gaussian and for which there is no exact FIM formula, we showed that the FSP-FIM is more accurate than traditional approaches (Figs. 4,5), which we validated by generating many independent data sets and comparing the inverse of the FSP-FIM to the variance in the MLE estimates (Figs. 3 and 6).

We showed that the choice of FIM analysis can lead to different optimal experiment designs (Fig. 5). For example, Figs. 5 and S3 show that the LNA-FIM can substantially overestimate the information of certain experiments compared to the full, correct information obtain using the FSP-FIM, which could mislead researchers to choose experiment designs that are much worse than they expect. In practice, overestimation of the Fisher information can have the further deleterious effect of overconfidence in poor parameter estimates, which can result in model bias and poor predictions as we observed recently in [26]. Furthermore, the LNA-FIM is not self-consistent, and often overestimates the information even compared to the ellipse found from sampling the MLE with the Gaussian likelihood function. On the other hand, we found that the SM-FIM under-estimated the information for the bursting gene model, but the amount of underestimation varied substantially with experimental conditions, which could cause researchers to reject otherwise informative experiments. In contrast to these moment-based approaches, the MLE sampling using the FSP approach always provided the best parameter estimates (Figs. 3 and S3), and the FSP-FIM was always consistent with the confidence intervals verified by sampling (Figs. 1, 3, 5, S1-S3), even for the case of nonlinear reaction propensities for which exact moments cannot be found (Figs. 6(a), and S4).

In our analysis of the non-linear toggle model, we allowed for the independent control of three experimental variables (Fig. 7a), and found experiments that optimize particular criteria of the FIM. Furthermore, we showed that other experiments very near to the optimal experiment in the design space can be significantly less informative than the optimal experiment (Figs. 7(b-e) and S6). Choosing between such similar experiment designs is non-trivial and would be difficult or impossible using intuition alone. On the other hand, we explored the effects of parameter uncertainty on FSP-FIM-based experiment design, and we found that parameter rankings are relatively robust to parameter uncertainty, even when the absolute value of the FSP-FIM is sensitive (Fig. 7).

We found that that the choice of optimal experiments depends on the number of experiments to be completed (Table 2). For example, the optimal set of two experiments may not contain the optimal single experiment. Sometimes, the FIM for a given experiment may be singular or nearly singular, indicating that the model under investigation is unidentifiable for the current parameterization and experiment design. In such a case, the FIM-eigenvectors corresponding to the near-zero eigenvalues indicate specific linear combinations of parameters that are poorly constrained (e.g., ‘sloppy’ directions [47]). If a second complementary experiment can shift the orientation of these sloppy vectors, then those parameters may yet be uncovered through combinations of multiple experiments. Alternatively, if a given combination of parameters remains unidentifiable for all admissible experiments, then these irrevocably sloppy directions may be used to reformulate the model into one that has a reduced set of fully identifiable parameters. We note that as one conducts new experiments and collects new data, parameter posteriors will need to be updated. As this occurs, optimal experiments may also need to be adjusted (e.g., through application of a Bayesian experiment design framework [48]), and future developments are needed to incorporate FSP-FIM computations within such iterative frameworks.

Our results show that the FSP-FIM performs better than previous approaches for gene regulation models with low molecule counts or nonlinear reaction rates. Previous studies have demonstrated many realistic systems for which such FSP can be used to identify and predict stochastic dynamics in numerous biological systems [2,6,19,26,29–32,49]. Each of these studies has used different experimental input signals, such as temporal salinity profiles [2,26], temperature [29], or chemical induction [19,30]. Modern optogenetic experiments promise to allow for even more robust and flexible control of input signals to control cellular behavior [7,50,51]. For such studies, the FSP-FIM could now be used to optimize these signals to achieve maximally informative experiments.

Like any other tool, the FSP-FIM also has its associated challenges. Our initial investigations focused on intrinsic stochastic fluctuations of small biochemical processes, and we used simulated data to verify our new computational tools. For models with large molecular counts of four or more species or with the addition of mechanisms to account for extrinsic variability, existing methods to solve the FSP-FIM will remain intractable until more efficient probability density based CME analyses can be developed to address such problems [52-56]. Until higher dimension CME approaches are developed, approximate moment-based experiment design methods, such as the SM-FIM and LNA-FIM, may remain the only available options to design experiments for large biochemical pathways. We also note that real experiments come with additional sources of noise, such as the errors or uncertainties associated with experimental measurements. For example, in smFISH data analysis, image processing settings give rise to variability in final RNA counts due to density dependent co-localization of RNA molecules. This measurement uncertainty may have a non-negligible effect on parameter inference, and future controlled experiments are needed to elucidate the degree to which such effects depend on optical imaging settings. Fortunately, such variabilities are easily incorporated within the framework of the FSP analysis. For example, previous work has used a simple linear transformation to adapt FSP analyses to include the effects of noisy GFP fluorescence emission and background autofluorescence when comparing integer-valued biochemical models to flow cytometry data in arbitrary continuous units of fluorescence [19]. Once adapted to take these transformations into account, the FSP-FIM could be used to design experiments to minimize the effects of measurement noise.

New experimental capabilities are creating an enormous potential to probe single-cell biological responses. These capabilities are making it ever more difficult to choose what species in the system to measure, whether to measure joint distributions (i.e. measure the RNA counts from multiple genes in the same cells) or marginal distributions (only measure RNA counts from a single gene at a time), or in what condition. Furthermore, different experiments have different costs, and the experimentalists must not only optimize their information about model parameters, but also consider the trade-off between collecting more data and the cost of a given experiment. By providing a new computational tool to iteratively improve models and design experiments for an important class of biological problems, the FSP-FIM will help to improve quantitative predictive modeling of gene expression.

## Materials and Methods

### Derivation of sensitivities for FSP models

The change of probability *p*(**x**_*l*_) with respect to small changes in parameter *θ*_*j*_ describes the sensitivity of the *l*^*th*^ state in the Markov process to the *j*^*th*^ parameter [33,57]. These local sensitivities can be calculated by transforming the linear ODEs describing the time evolution of the probabilities of the state space 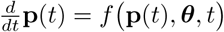 into a set of ODEs describing the time evolution of the sensitivities. Given an initial condition, the solution to the CME is:

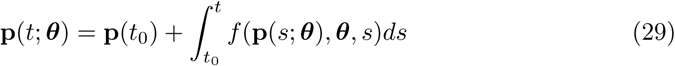

Taking partial derivatives with respect to ***θ***,

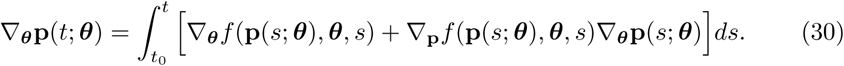

We can now describe the sensitivities **S** = ∇_***θ***_**p** as they evolve with time, by taking the time derivative of the equation above. For the FSP, the right-hand side *f* (**p**(*t*; ***θ***), ***θ****, t*) = **A**(***θ****, t*)**p**(*t*), and

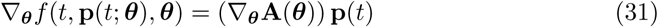

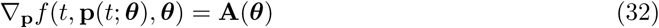

In many cases, including all models formulated using mass-action kinetics, the generator **A** can be written as a linear combination of the model parameters, i.e. **A** = Σ*θ*_*i*_**B**_*i*_, and the derivative with respect to the *i*^*th*^ parameter can be found,

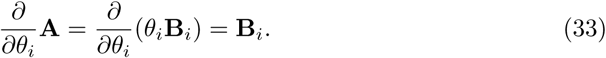

Using this notation, Eq. 30 is reduced to the set of linear ODEs for each parameter *θ*_*i*_,

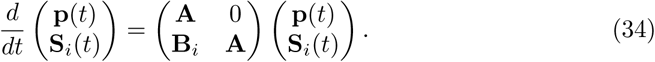

In practice, Eq. 34 can be computed in parallel for each parameter, and the computation of sensitivities for *K* parameters is equivalent to solving *K* sparse system of ODEs, each twice the size of the FSP computation.

### Moment-based FIM Approximations

Current state-of-the-art approaches for single-cell, single-molecule experiment design rely on computing moments of the CME. Such statistical moments may be computed exactly for systems with affine-linear propensities [42]. The uncentered moments of th CME, 𝔼{**x**^**m**^}, where **m** = [*m*_1_*, m*_2_*, …, m*_*N*_] is a vector of integers corresponding to the 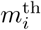 power of the *i*^th^ species in **x**, and the entire moment **x**^**m**^ is found according to the following formula:

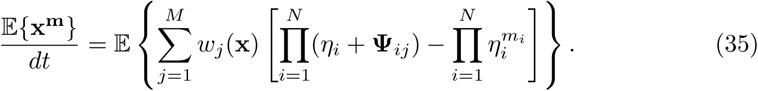

In the limit of large numbers of molecules reacting in a well-mixed solution, the linear noise approximation (LNA) may be applied to CME [25]. In such cases, molecule numbers are considered to be Gaussian, and the well-known Gaussian form of the FIM may be applied [8]. If the observed data is assumed to come from a multivariate Gaussian distribution with means ***µ***(*t*; ***θ***) = [*µ*_1_(*t*; ***θ***)*, µ*_2_(*t*; ***θ***)*, … µ*_*N*_ (*t*; ***θ***)]^*T*^ and covariance matrix **Σ**(*t*; ***θ***), such as those in Eqs. 35, the likelihood is given by:

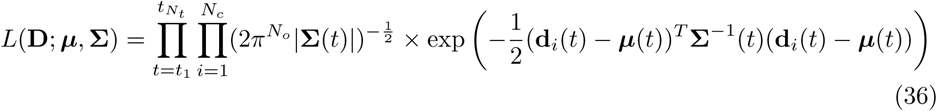

and the FIM is well-known [10, 11]

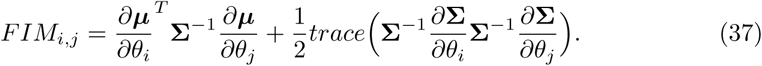

Another approach, developed in [9] is to use a likelihood function that takes the sample mean and sample variance to be jointly Gaussian, and thus requires the computation of up to the 4th moments in Eq. 35. In the supplement, we reproduce the formulae from [9] relevant to this study.

## Supporting information

## Supporting Information

### Supplemental text

Logarithmic parameterization of the FSP-FIM

Central Limit Theorem approximation

Generation and fitting of simulated data

Derivation of information for Gaussian fluctuations

Derivation of information for a Poisson distribution

## Supplemental figures

**Fig S1**. Optimal experiment design for the bursting gene expression model using the determinant of the FIM, D-optimality.

**Fig S2**. Optimal experiment design for the bursting gene expression model using E-optimality determined using the logarithmic parameterization of the FIM.

**Fig S3**. Verification of the optimal experiment design for the sampling period Δ*t* of a bursting gene expression model.

**Fig S4**. Verification of the FSP-FIM for the seven free parameters for the toggle model.

**Fig S5**. Sampled parameters for the uncertainty analysis of different experiment designs.

**Fig S6**. Different experiment designs’ effects on parameter uncertainty.

