## Supplementary material for "The finite state projection based Fisher information matrix approach to estimate information and optimize single-cell experiments"

#### Contents

**1 Logarithmic Parameterization of the FSP-FIM 2**

**2 Central Limit Theorem Approximation 2**

**3 Generation and fitting of simulated data 3**

**4 Derivation of information for Gaussian fluctuations 4**

**5 Derivation of information for a Poisson distribution 4**

### 1 Logarithmic Parameterization of the FSP-FIM

As noted in [1], in many biological systems, rates vary by several orders of magnitude. Therefore, it may be of interest to consider the relative change in parameter values and formulate the FIM in terms of the logarithm of the parameters,

$$\mathcal{I}(\log \boldsymbol{\theta}) = \mathbf{E}\Big[\Big(\nabla_{\log \boldsymbol{\theta}} \log p(\mathbf{X};\boldsymbol{\theta})\Big)^T \Big(\nabla_{\log \boldsymbol{\theta}} \log p(\mathbf{X};\boldsymbol{\theta})\Big)\Big] \tag{1}$$

The logarithmic parameterization carries through to the computation of the sensitivity matrix,

$$\nabla_{\boldsymbol{\theta}} \log p(x;\boldsymbol{\theta}) = \begin{pmatrix} \frac{1}{p_0}\frac{\partial p_0}{\partial \log \theta_1} & \frac{1}{p_0}\frac{\partial p_0}{\partial \log \theta_2} & \cdots & \frac{1}{p_0}\frac{\partial p_0}{\partial \log \theta_{N_p}} \\ \frac{1}{p_1}\frac{\partial p_1}{\partial \log \theta_1} & \frac{1}{p_1}\frac{\partial p_1}{\partial \log \theta_2} & \cdots & \frac{1}{p_1}\frac{\partial p_1}{\partial \log \theta_{N_p}} \\ \vdots & \vdots & \cdots & \vdots \\ \frac{1}{p_N}\frac{\partial p_N}{\partial \log \theta_1} & \frac{1}{p_N}\frac{\partial p_N}{\partial \log \theta_2} & \cdots & \frac{1}{p_N}\frac{\partial p_N}{\partial \log \theta_{N_p}} \end{pmatrix}. \tag{2}$$

Using the relationship $\frac{\partial f(x)}{\partial \log x} = x\frac{\partial f(x)}{\partial x}$, we can rewrite Eq. 2 as

$$\nabla_{\log \boldsymbol{\theta}} \log p(\mathbf{x};\boldsymbol{\theta}) = \mathbf{QS\Theta}, \tag{3}$$

where

$$\boldsymbol{\Theta} \equiv \begin{pmatrix} \theta_1 & 0 & \cdots & 0 \\ 0 & \theta_2 & \ddots & 0 \\ \vdots & \ddots & \ddots & \vdots \\ 0 & \cdots & \cdots & \theta_{N_p} \end{pmatrix}$$

and $\mathbf{Q} = diag\{\frac{1}{\mathbf{p}}\}$. Therefore, the logarithmic parameterization is easily found by multiplying the $i^{\text{th}}$ column in $\mathbf{S}$ by the corresponding parameter $\theta_i$. The log-FSP-FIM can then be computed:

$$\mathcal{I}(\log \boldsymbol{\theta})_{i,j} = N_c \sum_{k=1}^{N} \frac{\theta_i \theta_j}{p(\mathbf{x}_k;\boldsymbol{\theta})} \mathbf{s}_i^k \mathbf{s}_j^k = \theta_i \theta_j \mathcal{I}(\boldsymbol{\theta})_{ij}. \tag{4}$$

### 2 Central Limit Theorem Approximation

In [2], the authors suggest approximating the FIM for the sample mean and variance to be jointly Gaussian, i.e. the random vector $\mathbf{z} = [\boldsymbol{\mu}_{\mathbf{s}}, \boldsymbol{\Sigma}_s]^T$, $Z \sim \mathcal{N}(\mathbf{z}, \mathbf{C})$, and $\mathbf{C}$ is the covariance matrix:

$$\mathbf{C} = \begin{pmatrix} \mathbf{C}_{\boldsymbol{\mu}_s \boldsymbol{\mu}_s} & \mathbf{C}_{\boldsymbol{\mu}_s \boldsymbol{\Sigma}_s} \\ (\mathbf{C}_{\boldsymbol{\mu}_s \boldsymbol{\Sigma}_s})^T & \mathbf{C}_{\boldsymbol{\Sigma}_s \boldsymbol{\Sigma}_s} \end{pmatrix}. \tag{5}$$

The submatrices on the diagonal correspond to the variance of $\boldsymbol{\mu}_s$ and $\boldsymbol{\Sigma}_s$, and the off diagonal terms correspond to correlations between the sample means and variances.

In [2], they derive the elements of each of these matrices in terms of the moments of the underlying model distribution $p(\mathbf{x}|\boldsymbol{\theta})$ for models with one or two species.

For example, consider the variance/covariance of the sample mean is $\mathbf{C}_{\boldsymbol{\mu}_s\boldsymbol{\mu}_s}$, where we have the data matrix with $N$ measurements $\mathbf{X} = [\mathbf{x}_1^T\ \mathbf{x}_2^T \ldots \mathbf{x}_N^T]^T$, where each row in the matrix $\mathbf{X}$ corresponds to a different measurement. The sample mean $\bar{\mathbf{x}}$ can be written $\mathbf{1}^T\mathbf{X}/N$, where $\mathbf{1}$ is a column vector of ones of size $N$. Without loss of generality, let $\mathbf{E}[\mathbf{x}] = 0$, and

$$\mathbf{C}_{\boldsymbol{\mu}_s\boldsymbol{\mu}_s} = \frac{1}{N^2}\left(\mathbf{E}[\mathbf{1}^T\mathbf{X}\mathbf{X}^T\mathbf{1}] - \mathbf{E}[\mathbf{1}^T\mathbf{X}]\mathbf{E}[\mathbf{1}^T\mathbf{X}]^T\right) \tag{6}$$

$$= \frac{1}{N^2}\mathbf{E}[\mathbf{1}^T\mathbf{X}\mathbf{X}^T\mathbf{1}] = \frac{N}{N^2}\mathbf{E}[\mathbf{X}\mathbf{X}^T] \tag{7}$$

$$= \frac{1}{N}\boldsymbol{\Sigma}. \tag{8}$$

Similar procedures can be used find the rest of the $\mathbf{C}$, and are given in detail in the SI of [2].

In this paper, we compared the FIM for the bursting gene expression model, where only the sample moments were concerned. The formula for the sample-moments FIM for a single species using the sample mean and variance was derived in the SI of [2],

$$\mathcal{I}(\boldsymbol{\theta})_{i,j} = N\frac{\frac{\partial\langle x\rangle}{\partial\theta_i}\frac{\partial\langle x\rangle}{\partial\theta_j}}{\langle\tilde{x}^2\rangle} + N\frac{\left(\langle\tilde{x}^2\rangle\frac{\partial\langle\tilde{x}^2\rangle}{\partial\theta_i} - \frac{\partial\langle x\rangle}{\partial\theta_i}\langle\tilde{x}^3\rangle\right)\left(\langle\tilde{x}^2\rangle\frac{\partial\langle\tilde{x}^2\rangle}{\partial\theta_j} - \frac{\partial\langle x\rangle}{\partial\theta_j}\langle\tilde{x}^3\rangle\right)}{\langle\tilde{x}^2\rangle^2\left(\langle\tilde{x}^4\rangle - \langle\tilde{x}^2\rangle^2\right) - \langle\tilde{x}^2\rangle\langle\tilde{x}^3\rangle^2} + \mathcal{O}(1), \tag{9}$$

where $\tilde{x}$ refers to the centered moment, i.e. $\langle\tilde{x}^2\rangle = \langle(x - \langle x\rangle)^2\rangle$.

#### 3 Generation and fitting of simulated data

For the simulated studies of the bursting gene expression model and the toggle model, we generated 200 simulated data sets for each analysis by sampling the solution of the FSP solution at a reference parameter set $\boldsymbol{\theta}^*$. We used inverse transform sampling to generate 1,000 independent samples from the FSP solution at each time point, which correspond to RNA measurements in single cells. The FSP was solved to a precision of at least $10^{-4}$. For each data set, we found the parameters which maximize the likelihood of the data $\hat{\boldsymbol{\theta}}$ using a combination of global parameter search (via the Metropolis-Hastings algorithm) and local optimization routines (the Nelder-Mead and BFGS algorithms), implemented in the scientific Python library SciPy [3].

For the bursting gene expression model, three likelihood functions for each different approach were used. As described in the Methods section of the main text, the likelihood for a multivariate Gaussian is:

$$L(\mathbf{D};\boldsymbol{\mu},\boldsymbol{\Sigma}) = \prod_{t=t_1}^{t_{N_t}}\prod_{i=1}^{N_c}(2\pi^{N_o}|\boldsymbol{\Sigma}(t)|)^{-\frac{1}{2}} \tag{10}$$

$$\times \exp\Big[-\frac{1}{2}(\mathbf{d}_i(t) - \boldsymbol{\mu}(t,\boldsymbol{\theta}))^T\boldsymbol{\Sigma}^{-1}(t,\boldsymbol{\theta})(\mathbf{d}_i(t) - \boldsymbol{\mu}(t,\boldsymbol{\theta}))\Big]$$

For the LNA-based likelihood from [1], each $\mathbf{d}_i(t)$ corresponds to an RNA measurement in a single-cell, and $\boldsymbol{\mu}(t,\boldsymbol{\theta})$ and $\boldsymbol{\Sigma}(t,\boldsymbol{\theta})$ are both functions of the model parameters. We find the $\boldsymbol{\theta}$ which maximizes the logarithm of this function for each simulated data set. The sample-moments based likelihood instead compares the sample mean and sample variances as the vector $\mathbf{d}(t) = [\mu_s(t), \boldsymbol{\Sigma}_s(t)]$, for which there is only one value per data set (as all the data was used to find the sample mean and sample variance). In this case, $\boldsymbol{\mu}(t,\boldsymbol{\theta})$ is the model predicted average of the mean and variance, and $\mathbf{\Sigma}(t,\boldsymbol{\theta})$ is the model-predicted variance. For the one dimensional case of RNA in the bursting gene expression model,

$$\mathbf{\Sigma}(t,\boldsymbol{\theta}) = \begin{pmatrix} \langle \tilde{x}(t,\boldsymbol{\theta})^2 \rangle & \langle \tilde{x}(t,\boldsymbol{\theta})^3 \rangle \\ \langle \tilde{x}(t,\boldsymbol{\theta})^3 \rangle & \langle \tilde{x}(t,\boldsymbol{\theta})^4 \rangle - \frac{N-3}{N-1} \langle \tilde{x}(t,\boldsymbol{\theta})^2 \rangle^2 \end{pmatrix}. \tag{11}$$

The higher order moments of the model are computed according to Eq. 36 in the main text, and the likelihood is computed according to Eq. 37.

#### 4 Derivation of information for Gaussian fluctuations

The Gaussian distribution with mean and variance $\lambda$ is defined

$$f(x,\lambda) = \frac{1}{\sqrt{2\pi\lambda}} e^{\frac{(x-\lambda)^2}{2\lambda}}. \tag{12}$$

Computing the FIM for this Guassian requires finding the derivative of the log-density

$$\log f(x,\lambda) = -\frac{1}{2}\log 2\pi - \frac{1}{2}\log\lambda - \frac{1}{2}\left(\frac{x^2 - 2x\lambda + \lambda^2}{\lambda}\right) \tag{13}$$

with respect to $\lambda$,

$$\begin{aligned} \frac{\partial \log f(x,\lambda)}{\partial \lambda} &= -\frac{1}{2\lambda} - \frac{1}{2}(1 - \frac{x^2}{\lambda^2}) \\ &= -\frac{1}{2}(-\frac{x^2}{\lambda^2} + \frac{1}{\lambda} + 1) \end{aligned}$$

and squaring it:

$$\begin{aligned} \left(\frac{\partial \log f(x,\lambda)}{\partial \lambda}\right)^2 &= \frac{1}{4}\left(-\frac{x^2}{\lambda^2} + \frac{1}{\lambda} + 1\right)\left(-\frac{x^2}{\lambda^2} + \frac{1}{\lambda} + 1\right) \\ &= \frac{1}{4}\left(\frac{x^4}{\lambda^4} - \frac{2x^2}{\lambda^3} - \frac{2x^2}{\lambda^2} + \frac{1}{\lambda^2} + \frac{2}{\lambda} + 1\right). \end{aligned} \tag{14}$$

To take the expected value, we need the second and fourth moments of the normal distribution, which are $\lambda^2+\lambda$ for the second uncentered moment and $\lambda^4+6\lambda^3+3\lambda^2$ for the fourth uncentered moment. Thus, we have:

$$\begin{aligned} \mathbf{E}\Big[\Big(\frac{\partial \log f(x,\lambda)}{\partial \lambda}\Big)^2\Big] &= \frac{1}{4}\Big(\frac{\lambda^4 + 6\lambda^3 + 3\lambda^2}{\lambda^4} - \frac{2(\lambda^2+\lambda)}{\lambda^3} - \frac{2(\lambda^2+\lambda)}{\lambda^2} + \frac{1}{\lambda^2} + \frac{2}{\lambda} + 1\Big) \\ &= \frac{1}{4}\Big(\frac{4}{\lambda} + \frac{2}{\lambda^2}\Big) = \frac{1}{\lambda} + \frac{1}{2\lambda^2}. \end{aligned}$$

#### 5 Derivation of information for a Poisson distribution

The Poisson distribution is defined:

$$f(x,\lambda) = \frac{\lambda^x e^{-\lambda}}{x!}. \tag{15}$$

Again, by taking the log

$$\log f(x, \lambda) = x \log \lambda - \lambda - \log x! \tag{16}$$

Now, take the derivative with respect to $\lambda$

$$\frac{\partial \log f(x, \lambda)}{\partial \lambda} = \frac{x}{\lambda} - 1, \tag{17}$$

and squaring this term yields:

$$\Big(\frac{\partial \log f(x, \lambda)}{\partial \lambda}\Big)^2 = \frac{x^2}{\lambda^2} - \frac{2x}{\lambda} + 1. \tag{18}$$

As the FIM is the expected value of this quantity, and the mean and variance of the Poisson distribution are given by $\lambda$,

$$\begin{aligned} \mathbf{E}\Big[\Big(\frac{\partial \log f(x, \lambda)}{\partial \lambda}\Big)^2\Big] &= \mathbf{E}\Big[\frac{x^2}{\lambda^2}\Big] - \mathbf{E}\Big[\frac{2x}{\lambda}\Big] + 1 \\ &= \frac{\lambda^2 + \lambda}{\lambda^2} - 2 + 1 \\ &= \frac{1}{\lambda}. \end{aligned} \tag{19}$$

| | $\log b_y$ | $\log b_x$ | $\log k_y$ | $\log k_x$ | $\log \alpha_{xy}$ | $\log \alpha_{yx}$ | $\log \gamma_x$ |
|---|---|---|---|---|---|---|---|
| $\log b_y$ | 0.673 | 0.130 | -0.017 | 0.194 | 0.165 | 0.039 | 0.135 |
| $\log b_x$ | 0.130 | 0.182 | 0.010 | 0.177 | 0.082 | -0.011 | 0.184 |
| $\log k_y$ | -0.017 | 0.010 | 0.004 | 0.007 | 0.007 | 0.006 | 0.009 |
| $\log k_x$ | 0.194 | 0.176 | 0.007 | 0.212 | 0.093 | 0.041 | 0.177 |
| $\log \alpha_{xy}$ | 0.165 | 0.082 | 0.007 | 0.093 | 0.084 | 0.038 | 0.08 |
| $\log \alpha_{yx}$ | 0.038 | -0.011 | 0.006 | 0.041 | 0.038 | 0.160 | -0.023 |
| $\log \gamma_x$ | 0.135 | 0.184 | 0.009 | 0.177 | 0.080 | -0.023 | 0.187 |

Table 1: Variance and covariance of the log of each parameter for the toggle model for prior uncertainty in the toggle model. This covariance was chosen according to the inverse of the logarithmic parameterized FIM evaluated for an experiment with 0 UV, $t = [1, 4, 8]$ hr, and 100 measurements at each time point.

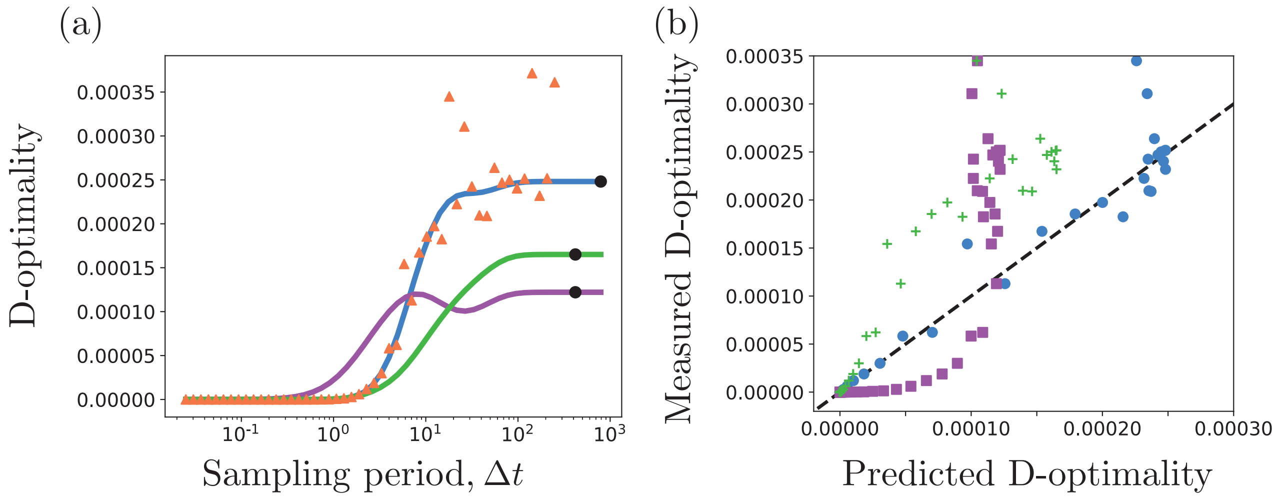

Figure S1: Optimal experiment design for the bursting gene expression model using the determinant of the FIM, D-optimality. (a) The D-optimality criteria for the FSP-FIM (blue), LNA-FIM (purple) and SM-FIM (green) for different sampling periods $\Delta t$. Orange triangles represent the D-optimality confirmed using 200 simulated data sets for each potential sampling period. Optimal sampling periods are given by black circles. (b) Comparison of the FSP-FIM at the the reference parameter set (x-axis) and the observed information (y-axis) for various sampling periods using the FSP-FIM (blue circles), LNA-FIM (purple squares), and SM-FIM (green crosses). Kinetic parameters are $k_{on} = 0.05$ min$^{-1}$, $k_{off} = 0.15$ min$^{-1}$, $k_r = 5$ molecules/min, and $\gamma = 0.05$ min$^{-1}$.

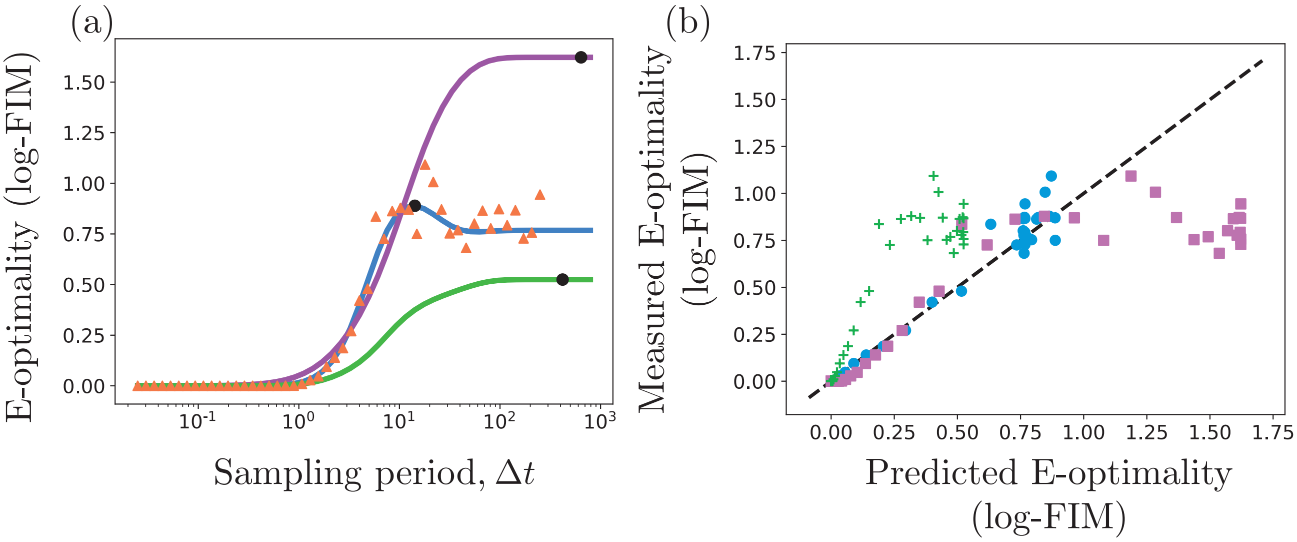

Figure S2: Optimal experiment design for the bursting gene expression model using E-optimality determined using the logarithmic parameterization of the FIM. The FSP-FIM (blue), LNA-FIM (purple) and SM-FIM (green) are shown for different sampling periods $\Delta t$. Orange triangles represent the E-optimality confirmed using 200 simulated data sets for each potential sampling period, where the optimal sampling periods are given by black circles.

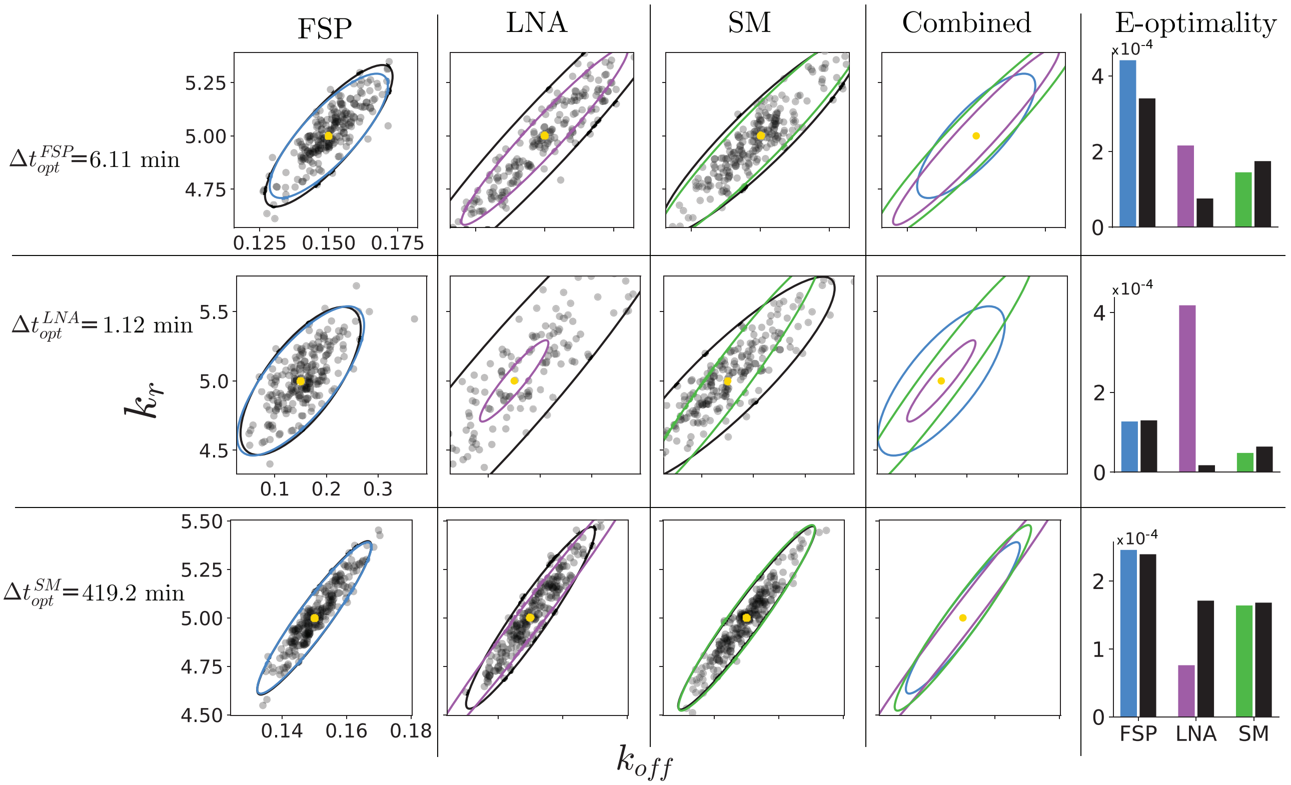

Figure S3: Verification of the optimal experiment design for the sampling period $\Delta t$ of a bursting gene expression model. For each optimal sampling period, we simulated 200 data sets and find the MLE estimates for each data set (black circles) using each likelihood function (FSP, LNA and sample moments). The top row uses a $\Delta t$ according to the optimal sampling period found using the FSP-FIM, the middle row uses the optimal $\Delta t$ found using the LNA-FIM, and the bottom row using the SM-FIM, using the regular parameterization of the FIM (i.e. the black circles in Fig. 5).

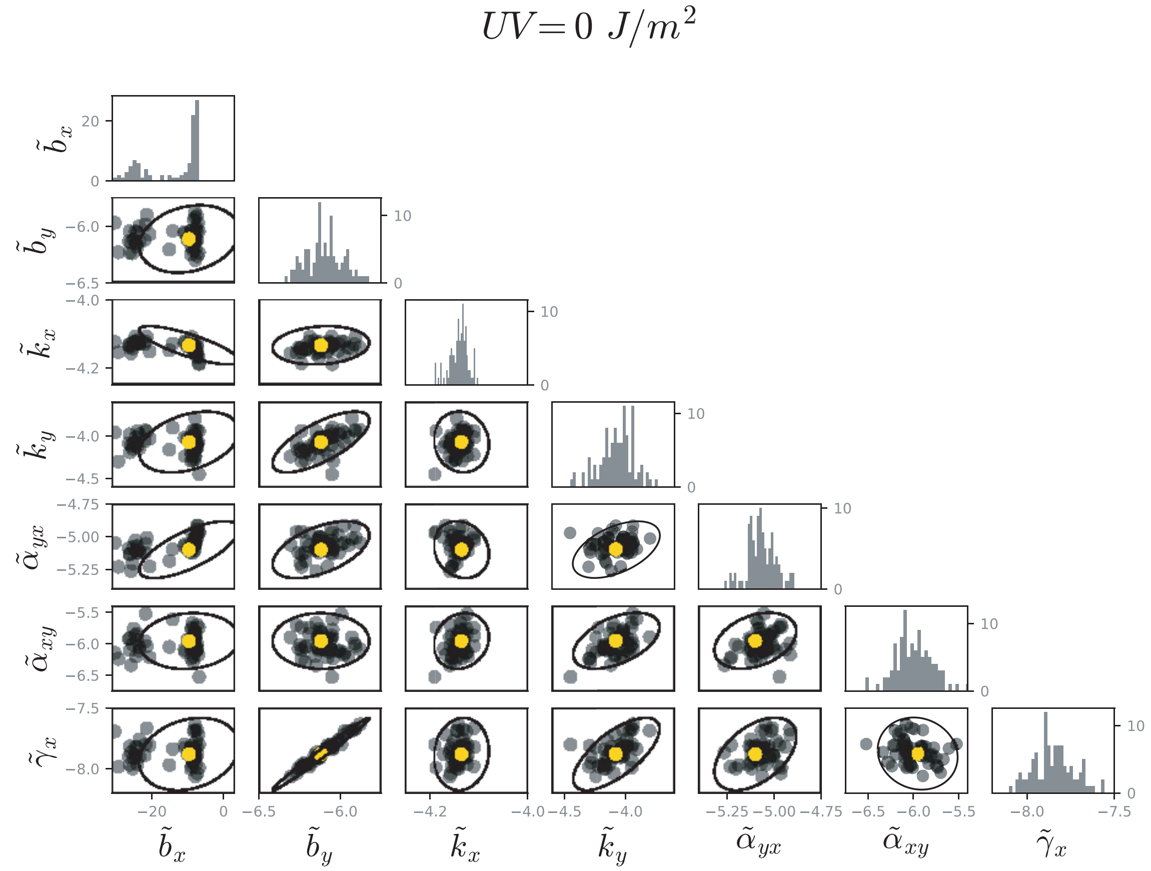

Figure S4: Verification of the FSP-FIM for the seven free parameters for the toggle model. Each black circle corresponds to the logarithm of an MLE estimate, $\log \hat{\boldsymbol{\theta}}$ for 100 different simulated data sets. The gold circle corresponds to the reference parameter set, $\log \boldsymbol{\theta}^*$. The 95% ellipse corresponding to the log-FSP-FIM is shown in black. The tilde corresponds to the log of each parameter, i.e. $\tilde{b}_x = \log b_x$.

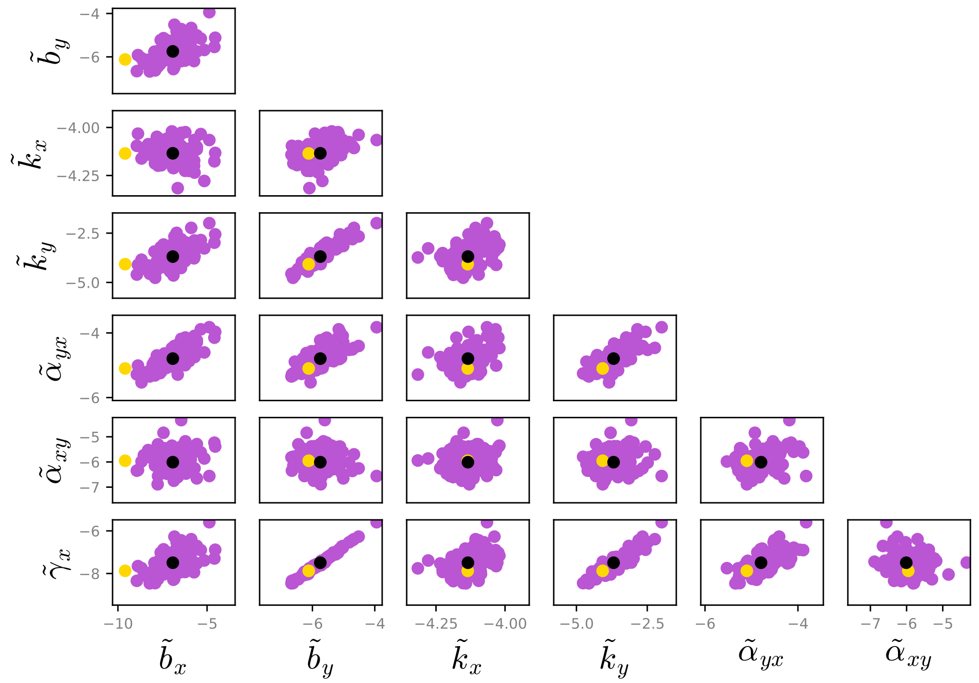

Figure S5: Parameters were sampled 100 times from a log-normal distribution evaluated about the reference parameter set $\hat{\boldsymbol{\theta}}_0$, shown in black. The covariance of this distribution was chosen according to the inverse of the FIM evaluated for an experiment with 0 UV, $t = [1, 4, 8]$ hr, and 100 measurements at each time point. For reference, the gold parameters are the 'true' parameters for the model. The tilde corresponds to the log of each parameter, i.e. $\tilde{b}_x = \log b_x$.

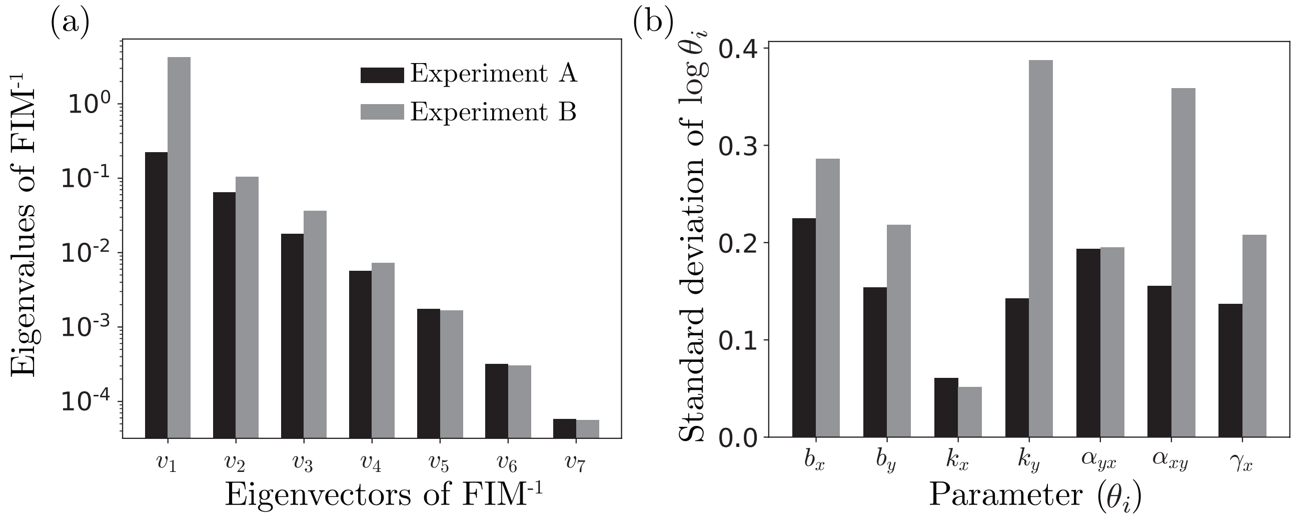

Figure S6: (a) The eigenvalues of the inverse of the Fisher information (i.e. the CRB) for the two experiments in Fig. 7(c) in the main text. Note that the y-axis is on a logarithmic scale. Lower values correspond to lower parameter uncertainty. (b) The effect of Experiment A and B on standard deviations of $\log \theta_i$.
